# *Coxiella burnetii* small RNA 12 binds CsrA regulatory protein and transcripts for the CvpD type IV effector, regulates pyrimidine and methionine metabolism, and is necessary for optimal intracellular growth and vacuole formation during infection

**DOI:** 10.1101/679134

**Authors:** Shaun Wachter, Matteo Bonazzi, Kyle Shifflett, Abraham S. Moses, Rahul Raghavan, Michael F. Minnick

## Abstract

*Coxiella burnetii* is an obligate intracellular gammaproteobacterium and zoonotic agent of Q fever. We previously identified 15 small non-coding RNAs (sRNAs) of *C. burnetii*. One of them, named CbsR12 (***<u>C</u>****oxiella **<u>b</u>**urnetii* **<u>s</u>**mall **<u>R</u>**NA **<u>12</u>**) is highly expressed during growth in axenic medium and becomes even more dominant during infection of cultured mammalian cells. Secondary structure predictions of CbsR12 revealed four putative CsrA-binding sites in single-stranded segments of stem loops with consensus AGGA/ANGGA motifs. From this foundation, we determined that CbsR12 binds to recombinant *C. burnetii* CsrA-2, but not CsrA-1, proteins *in vitro*. Moreover, through a combination of *in vitro* and *in vivo* assays, we identified several in *trans* mRNA targets of CbsR12. Of these, we determined that CbsR12 binds to and upregulates translation of *carA* transcripts coding for carbamoyl phosphate synthetase A; an enzyme that catalyzes the first step of pyrimidine biosynthesis. In addition, CbsR12 binds and downregulates translation of *metK* transcripts coding for S-adenosyl methionine (SAM) synthase, a component of the methionine cycle. Furthermore, we found that CbsR12 binds to and downregulates the quantity of *cvpD* transcripts, coding for a type IVB effector protein, *in vitro* and *in vivo*. Finally, we found that CbsR12 is necessary for full expansion of *Coxiella*-containing vacuoles (CCVs) and affects bacterial growth rates in a dose-dependent manner in the early phase of infecting THP-1 cells. This is the first detailed characterization of a *trans*-acting sRNA of *C. burnetii* and the first example of a bacterial sRNA that regulates both CarA and MetK expression. CbsR12 is also one of only a few identified *trans-*acting sRNAs that interacts with CsrA. Results illustrate the importance of sRNA-mediated regulation in establishment of the intracellular CCV niche.

**Author summary:** *C. burnetii* is an obligate intracellular bacterial pathogen that is transmitted to humans from animal reservoirs. Upon inhalation of aerosolized *C. burnetii*, the agent is phagocytosed by macrophages in the lung. The pathogen subverts macrophage-mediated degradation and resides in a large, intracellular, acidic vacuole, termed the *Coxiella*-containing vacuole (CCV). Small RNAs (sRNAs) are not translated into proteins. Instead, they target mRNAs in order to up- or down-regulate their stability and translation. Alternatively, some sRNAs bind to regulatory proteins and serve as “sponges” that effectively sequester the proteins and inhibit their function. *C. burnetii*’s CbsR12 sRNA is highly expressed during infection in order to expand the CCV, and it works by a variety of mechanisms, including: 1) directly regulating transcripts of several metabolic genes that aid in bacterial replication, 2) binding to and regulating transcripts of a type IV effector protein that aids in infection, and 3) indirectly regulating an unknown number of genes by binding to a homolog of the global regulatory protein, CsrA. CbsR12 represents one of only a few sRNAs known to bind and sequester CsrA while also directly regulating mRNAs.

## Introduction

*Coxiella burnetii* is a Gram-negative, obligate intracellular bacterium and the etiological agent of Q fever in humans. Q fever most often manifests as an acute, flu-like illness, which in rare cases progresses to potentially life-threatening endocarditis [1]. *C. burnetii* undergoes a biphasic life cycle in which it alternates between a metabolically-active, replicative large-cell variant (LCV) and a non-replicative, spore-like small-cell variant (SCV) [2]. Upon aerosol transmission of SCVs to a mammalian host, *C. burnetii* is primarily endocytosed by alveolar macrophages, after which it survives acidification of the host phagolysosome and metamorphoses to LCVs. *C. burnetii* then utilizes the fusion of its *Coxiella*-containing vacuole (CCV) with lysosomes and autophagosomes in order to expand the intracellular niche [3, 4]. CCV expansion is dependent on *C. burnetii* protein synthesis, but independent of *C. burnetii* replication, so expansion of the CCV is facilitated by a repertoire of Dot/Icm effector proteins secreted by a Type IV-B secretion system (T4BSS) [5, 6]. Many Dot/Icm substrates have been identified in recent years [7] and shown to modulate the host inflammasome [8], influence autophagosomal/lysosomal fusion with the CCV by various mechanisms [9–16], and regulate the host transcriptome after localizing to the nucleus [17, 18]. Little is known about regulation of the *C. burnetii* T4BSS, although the PmrA response regulator has been shown to enhance expression of the T4BSS apparatus as well as certain Dot/Icm substrates [19].

Bacterial sRNAs are small (<500 nts) transcripts that do not code for functional proteins. Instead, they serve as *cis-* and/or *trans-*acting regulators through a variety of mechanisms [reviewed in [20]]. For example, *cis*-acting sRNAs are often coded antisense to a functional gene target. Upon transcription, the sRNA binds to the mRNA with perfect complementarity, usually culminating in ribonuclease degradation of the target. This effectively limits the free mRNA molecules available for translation [reviewed in [21]]. Alternatively, *trans*-acting sRNAs are often coded in distant intergenic regions (IGRs) and bind to a variety of mRNAs through a more limited base-pairing mechanism involving a seed region of around 7 - 12 nts. Many *trans*-acting sRNAs have been discovered in bacteria since *Escherichia coli* MicF was first described in 1984 [22]. These regulatory RNAs have been implicated in a variety of processes, including virulence [23], global regulation of transcription [24], iron homeostasis [25], protein degradation [26], and stress response [27, 28].

Typically, *trans*-acting sRNAs require assistance in “finding” their respective mRNA targets. In most bacteria, this is accomplished by the RNA chaperone Hfq, which binds to both sRNAs and mRNAs and plays the role of a molecular matchmaker [reviewed in [32]]. Hfq is not obligatory, however. For example, *Staphylococcus aureus* has several sRNAs but does not require Hfq protein for their activities [33]. Similarly, *C. burnetii* does not have a readily apparent *hfq* gene, although this doesn’t rule out the possibility of an atypical Hfq or some other novel RNA chaperone. For instance, *C. burnetii* codes for two homologs (CsrA-1, CsrA-2) of the RNA-binding protein CsrA (RsmA), which has been shown to regulate metabolism, biofilm formation, and Type 4 secretion in other bacteria [34–36]. CsrA is regulated by CsrA-binding sRNAs, termed CsrB/C (RsmY/Z). Classical CsrB/C sRNAs consist of a series of stem-loops containing exposed AGGA or ANGGA motifs that bind and sequester CsrA, effectively limiting its mRNA regulatory capabilities [37]. Some RsmY/Z sRNAs, however, differ in the number of stem-loop regions containing CsrA-binding sites, and can harbor far fewer motifs than the classical CsrB/C *E. coli* counterparts [38, 39]. The CsrA regulatory cascade has not been studied in *C. burnetii*, in large part due to the absence of readily-discernible RsmY/Z sRNAs, although the CsrA regulon in *Legionella pneumophila*, a close relative of *C. burnetii*, has been extensively studied [40, 41].

A previous study by our group revealed 15 novel *C. burnetii* sRNAs that were differentially expressed either in LCVs vs. SCVs, or in host cell infections vs. growth in ACCM-2 medium [29, 30]. Among these, CbsR12 was found to be markedly upregulated in the intracellular niche as compared to ACCM-2. Northern blots also showed that CbsR12 was upregulated in SCVs vs. LCVs in ACCM-2, and revealed two distinct sizes of the sRNA, suggesting that either an alternative transcription start site (TSS) or ribonuclease processing of the sRNA was responsible. In a subsequent study, CbsR3 and CbsR13 were found to originate from transcribed loci of a selfish genetic element, termed QMITE1 [31]. However, despite the identification and verification of several CbsRs, none has been functionally characterized, to date.

In this study, we describe functions of a highly expressed, infection-specific sRNA of *C. burnetii*, termed CbsR12. Our analyses show that CbsR12 binds to CsrA-2, but not CsrA-1 *in vitro*. We also establish that CbsR12 binds to and up-and down-regulates *carA* and *metK* transcripts, respectively, in *trans*. The bacterial *carA* gene codes for carbamoyl-phosphate synthetase (small) subunit A (CarA) which forms a heterodimer with carbamoyl-phosphate synthetase (large) subunit B (CarB). The CarAB complex catalyzes the first step in pyrimidine biosynthesis and is involved in arginine biosynthesis in some bacteria [42]. The bacterial *metK* gene codes for S-adenosyl L-methionine (SAM) synthase, an enzyme responsible for catalyzing the production of SAM; the major donor of methyl groups during metabolism in prokaryotic cells. As a methyl donor, SAM affects DNA methylation and thus global transcription [43]. It has also been implicated in virulence, being necessary for the production of N-acyl homoserine lactones involved in bacterial quorum sensing [reviewed in [44]]. We also implicate CbsR12 in the expansion of the *C. burnetii* vacuole, as the size of CCVs is directly correlated with levels of the sRNA. Furthermore, we find that CbsR12 binds *ahcY* transcripts, another component of the methionine cycle, *in vivo*, and *cvpD*, a T4BSS effector protein necessary for proper CCV development, *in vitro* and *in vivo* [11]. Overall, this study highlights CbsR12 as a crucial component in the early stages of *Coxiella* infection.

## Results

### CbsRh12 is a dominant non-rRNA/tRNA/tmRNA transcript during *C. burnetii* infection of Vero and THP-1 cells

CbsR12 was first described as a highly expressed, infection-specific sRNA that was upregulated in SCVs compared to LCVs when analyzed by Northern blots [29]. The impetus for our study came when we analyzed previous RNA-Seq data (SRP041556) by converting raw read data into transcripts per million (TPM), a normalized measure of gene expression [45]. These results showed that CbsR12 was the dominant non-tmRNA transcript in both LCVs and SCVs during *C. burnetii* infection of African green monkey kidney epithelial cells (Veros). Additional data from LCVs obtained from a *C. burnetii* infection of monocytic THP-1 cells corroborates the observation that CbsR12 is a dominant transcript during infection (Rahul Raghavan, unpublished results). Moreover, we were surprised to find that CbsR12 was more abundant in LCVs, not SCVs (Table 1).

**Table 1.**
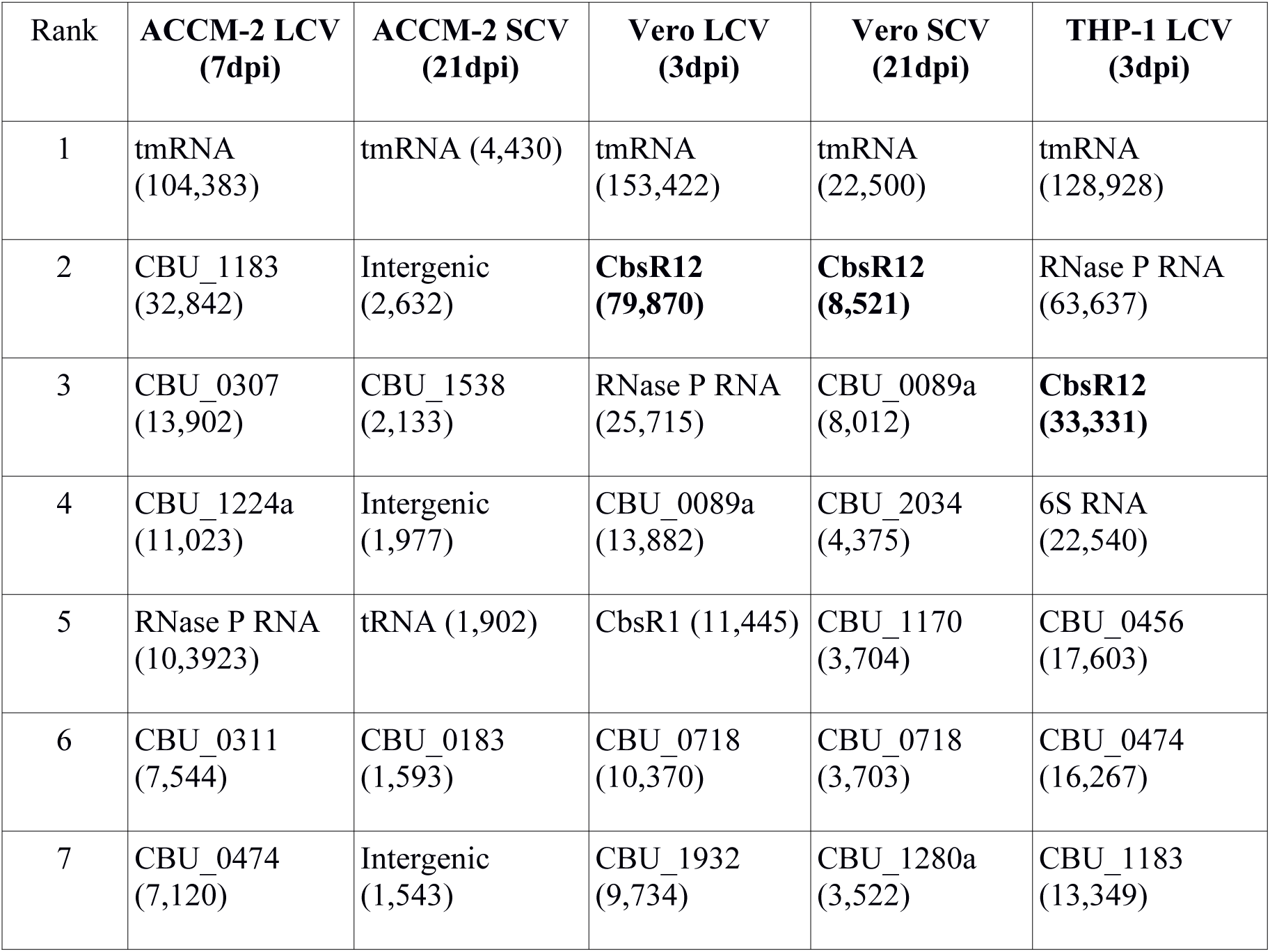
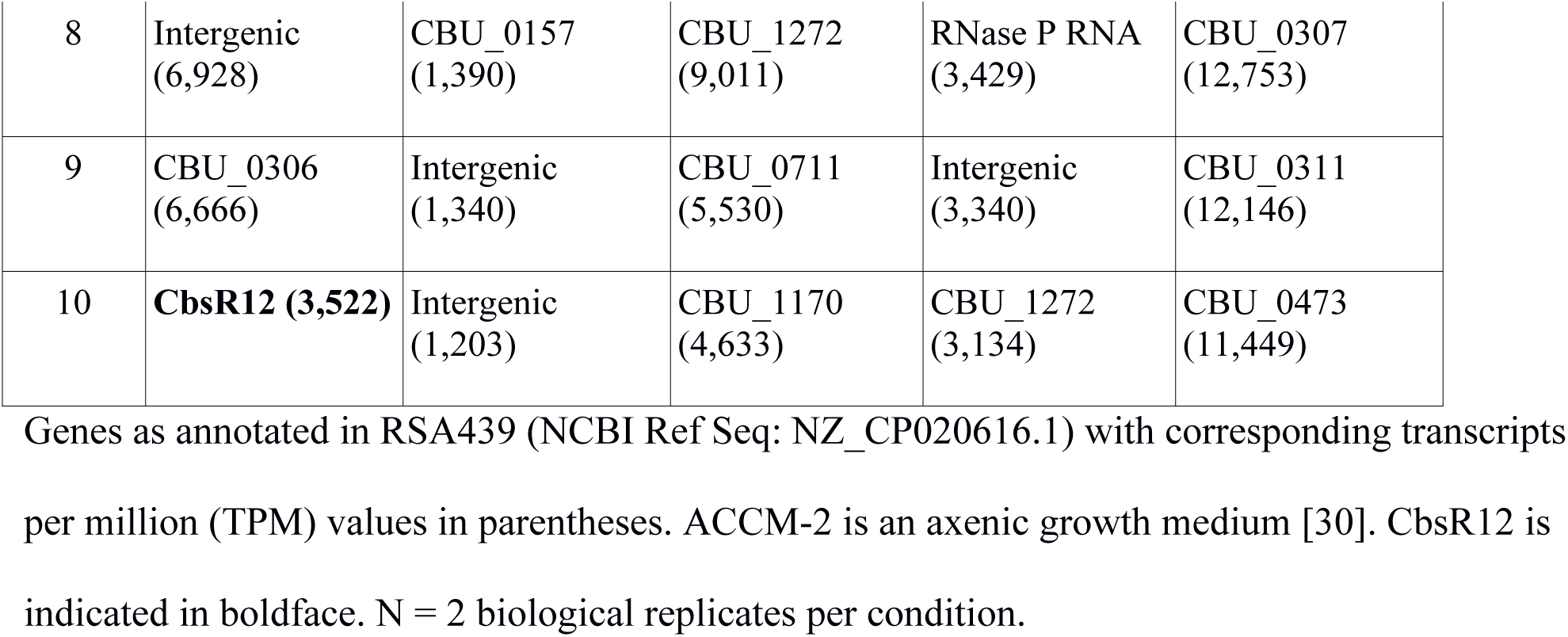
Top ten expressed genes across various *C. burnetii* growth conditions, ranked.

| Rank | ACCM-2 LCV<br>(7dpi) | ACCM-2 SCV<br>(21dpi) | Vero LCV<br>(3dpi) | Vero SCV<br>(21dpi) | THP-1 LCV<br>(3dpi) |
| --- | --- | --- | --- | --- | --- |
| 1 | tmRNA<br>(104,383) | tmRNA (4,430) | tmRNA<br>(153,422) | tmRNA<br>(22,500) | tmRNA<br>(128,928) |
| 2 | CBU_1183<br>(32,842) | Intergenic<br>(2,632) | **CbsR12**<br>**(79,870)** | **CbsR12**<br>**(8,521)** | RNase P RNA<br>(63,637) |
| 3 | CBU_0307<br>(13,902) | CBU_1538<br>(2,133) | RNase P RNA<br>(25,715) | CBU_0089a<br>(8,012) | **CbsR12**<br>**(33,331)** |
| 4 | CBU_1224a<br>(11,023) | Intergenic<br>(1,977) | CBU_0089a<br>(13,882) | CBU_2034<br>(4,375) | 6S RNA<br>(22,540) |
| 5 | RNase P RNA<br>(10,3923) | tRNA (1,902) | CbsR1 (11,445) | CBU_1170<br>(3,704) | CBU_0456<br>(17,603) |
| 6 | CBU_0311<br>(7,544) | CBU_0183<br>(1,593) | CBU_0718<br>(10,370) | CBU_0718<br>(3,703) | CBU_0474<br>(16,267) |
| 7 | CBU_0474<br>(7,120) | Intergenic<br>(1,543) | CBU_1932<br>(9,734) | CBU_1280a<br>(3,522) | CBU_1183<br>(13,349) |
| 8 | Intergenic<br>(6,928) | CBU_0157<br>(1,390) | CBU_1272<br>(9,011) | RNase P RNA<br>(3,429) | CBU_0307<br>(12,753) |
| 9 | CBU_0306<br>(6,666) | Intergenic<br>(1,340) | CBU_0711<br>(5,530) | Intergenic<br>(3,340) | CBU_0311<br>(12,146) |
| 10 | **CbsR12 (3,522)** | Intergenic<br>(1,203) | CBU_1170<br>(4,633) | CBU_1272<br>(3,134) | CBU_0473<br>(11,449) |

Genes as annotated in RSA439 (NCBI Ref Seq: NZ\_CP020616.1) with corresponding transcripts per million (TPM) values in parentheses. ACCM-2 is an axenic growth medium [30]. CbsR12 is indicated in boldface. N = 2 biological replicates per condition.

### CbsR12 is processed by ribonuclease III *in vitro*

Previous Northern blot analyses on sRNAs of *C. burnetii* showed that CbsR12 produced two distinct bands of approximately 170 and 50 nucleotides, regardless of growth conditions or developmental stage [29]. We therefore wished to determine whether these bands arose from alternative TSSs for the *cbsR12* gene or from ribonuclease III (RNase III) processing of the full-length CbsR12 transcript. We first utilized 5’ rapid amplification of cDNA ends (RACE) to determine the TSS of CbsR12. This experiment revealed the full-size CbsR12 (∼200 nucleotides) as expected, but also indicated that two potential alternative TSSs existed ∼110 nucleotides upstream of the *cbsR12* gene’s Rho-independent terminator (**S1A Fig**). To determine whether the TSSs were generated by RNase-mediated cleavage, we treated *in vitro-*transcribed CbsR12 with recombinant *C. burnetii* RNase III [40] and a commercially-available *E. coli* RNase III (New England BioLabs). Results showed that CbsR12 was processed by both kinds of RNase III into two RNA fragments, and the resulting sizes closely resembled those observed in the previous Northern blot analysis (**S1B Fig**) [29]. These results strongly suggest that the two sites are not alternative TSSs but are instead generated by RNase III processing.

### CbsR12 binds to *C. burnetii* recombinant CsrA-2, but not CsrA-1, *in vitro*

The *in silico*-predicted secondary structure of CbsR12 also revealed conserved single-strand sequence motifs among the various stem-loop structures (**S1A Fig**). This motif, AGGA/ANGGA, corresponds exactly to the conserved CsrA-binding motif of many bacteria [34, 36, 38]. *C. burnetii* contains two annotated and distinct types of CsrA (termed CsrA-1 and CsrA-2) that share 65% primary sequence identity (data not shown). To test the functionality of these domains, we employed an *in vitro* binding assay and a RNA-protein electrophoretic mobility shift assay (EMSA) to determine if CbsR12 binds to natively-purified recombinant CsrA-1 (rCsrA1) and/or CsrA-2 (rCsrA-2). EMSA results clearly showed that CbsR12 binds to rCsrA-2, but not rCsrA-1, *in vitro* (Fig 1). Furthermore, the K_D_ for rCsrA-2 was determined to be 130 nM; consistent with published values for CsrA-binding sRNAs [47].

**Fig 1.**
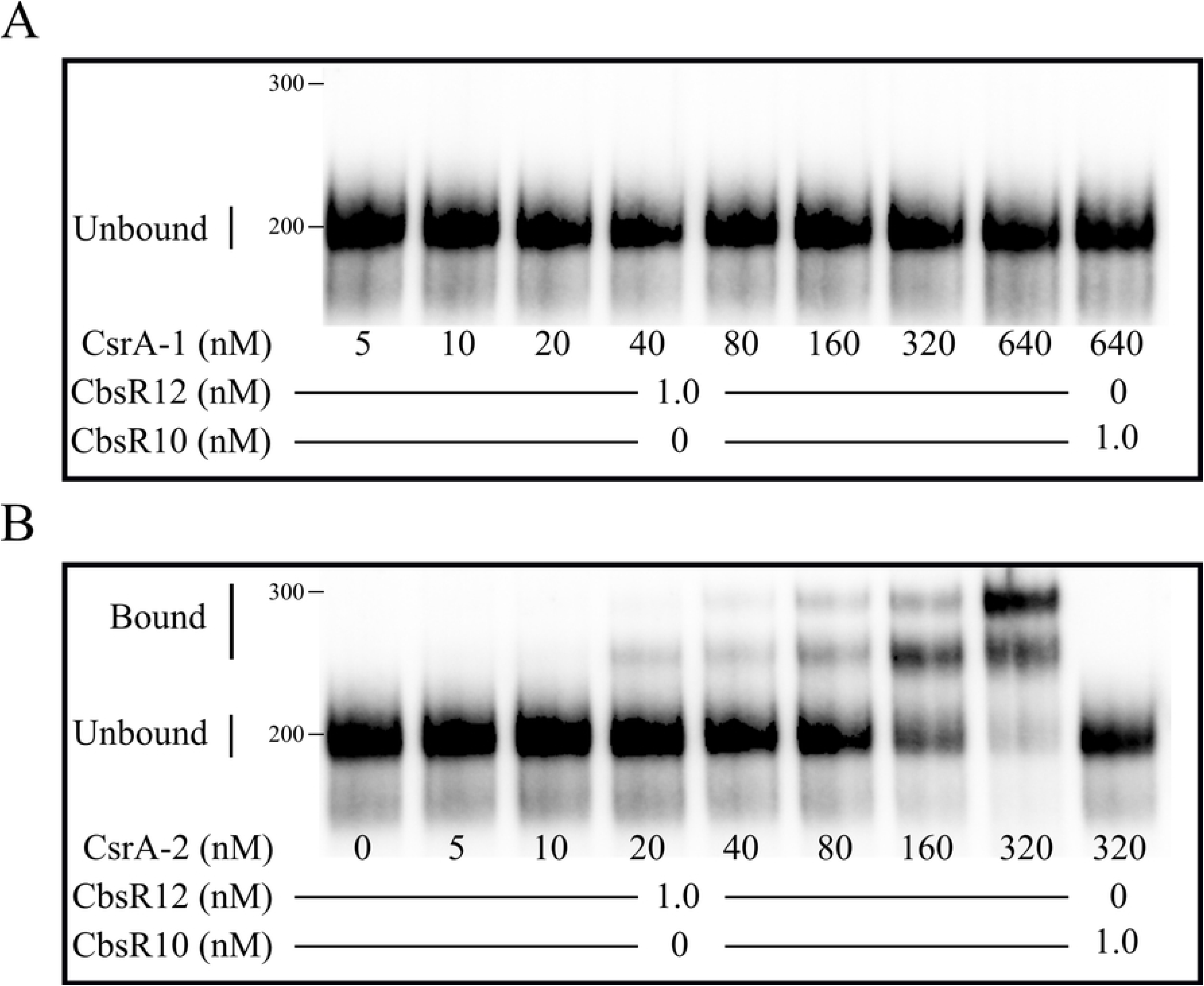
CbsR12 binds to CsrA-2, but not CsrA-1, protein *in vitro*. EMSAs showing RNA-protein interactions between biotin-labeled, *in vitro*-transcribed CbsR12 (0 or 1 nM) and increasing concentrations of purified, recombinant *C. burnetii* CsrA-1 (**A**) or CsrA-2 (**B**). CbsR10 (at 0 or 1 nM) is included as a negative control, as the sRNA contains only a single discernible CsrA-binding motif.

### A *cbsR12* mutant shows prolonged lag phase in axenic media

A *cbsR12* mutant (strain *Tn327*), as well as an otherwise isogenic transposon insertional control strain (*Tn1832*) of *C. burnetii*, were generated using a Himar1-based transposon system [14]. The location of the transposon insertion of strain *Tn327* is shown in **S2A Fig**. We also constructed a transposon-directed complement of strain *Tn327* (*Tn327-Comp*) containing the wild-type *cbsR12* gene plus ∼100 bp of 5’ and 3’ flanking sequences, to include any potential transcriptional regulator element(s) that could influence expression of *cbsR12*. PCR was used to confirm the transposon insertions in the *Tn327* and *Tn327-Comp* strains (**S2B Fig**).

Next, we conducted growth curve analyses of the *Tn1832*, *Tn327*, and *Tn327-Comp* strains grown axenically in ACCM-2 (Fig 2A) and assayed CbsR12 expression at incremental time points from the LCV stage (approximately 1-6 days) through the SCV stage (7+ days) by quantitative real-time PCR (qRT-PCR) (Fig 2B). Growth curve results showed that *Tn327* displayed a prolonged lag phase from 1-3 days post-inoculation that was not observed in *Tn1832* or *Tn327-Comp* strains (Fig 2A). Following lag phase, *Tn327* grew at a slightly increased rate relative to the other strains (days 6-9 post-infection), but failed to reach cell numbers seen in the other two strains throughout the assay. The “wild-type” (*Tn1832*) and complemented (*Tn327-Comp*) strains produced essentially indistinguishable growth curves. The qRT-PCR results showed that the *Tn327* insertion completely abrogated CbsR12 expression (Fig. 2B). The results also confirmed CbsR12’s increased expression in LCVs compared to SCVs as the copies of CbsR12 per *C. burnetii* genome were highest at 3 days post-inoculation.

**Fig 2.**
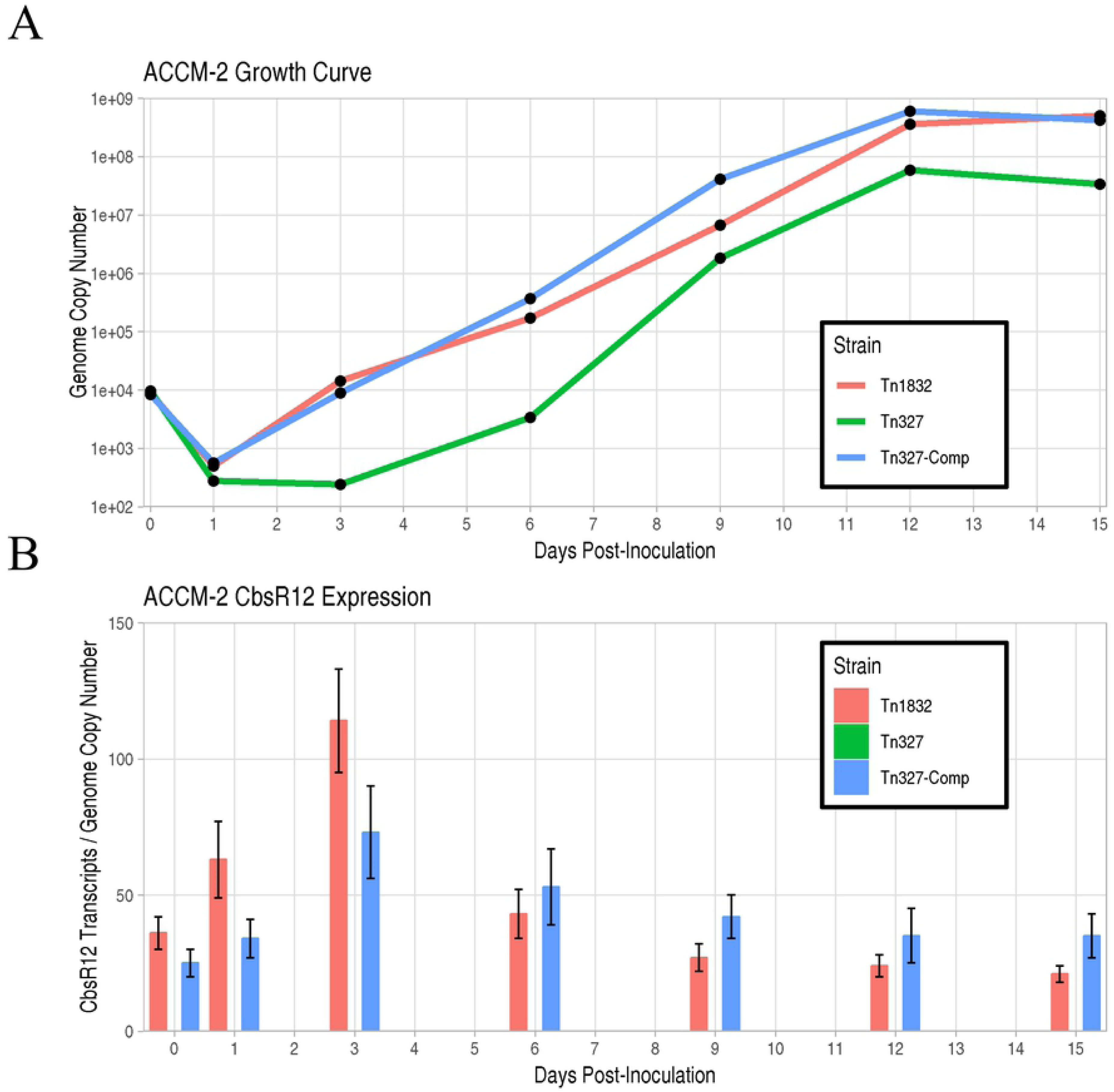
CbsR12 expression and growth effects on *C. burnetii* grown in ACCM-2. (**A**). Growth curves for *Tn1832*, *Tn327*, and *Tn327-Comp C. burnetii* strains in ACCM-2 as determined by qPCR. The 0dpi time point refers to the inoculum. Data are representative of three independent experiments with consistent and indistinguishable results. (**B**). CbsR12 expression over time for *Tn1832*, *Tn327*, and *Tn327-Comp C. burnetii* strains grown in ACCM-2 as determined by qRT-PCR. Values represent the means ± standard error of means (SEM) of three independent determinations.

### CbsR12 impacts intracellular replication of *C. burnetii*

*C. burnetii* typically infects alveolar macrophages during human infection. We therefore infected a differentiated human monocyte cell line (THP-1) with the *Tn1832*, *Tn327*, and *Tn327-Comp* strains. Comparative growth curves show that strain *Tn327* has a slower growth rate in exponential phase (1-3dpi) as compared to the two other strains, and never attains the bacterial cell numbers seen in infections with *Tn1832* or *Tn327-Comp* (Fig 3A). Furthermore, CbsR12 expression in THP-1s correlated with replication efficiency of the individual strain. For example, expression of CbsR12 in the *Tn327* and *Tn1832-Comp* strains increases between 1dpi and 3dpi, and CbsR12 levels directly correlates to growth rates of the respective strains between these two time points. However, we observed a dysregulation of CbsR12 in *Tn327-Comp* infecting THP-1 cells that was strikingly different than what was seen during axenic growth. Specifically, in THP-1s, we observed a maintenance of CbsR12 expression throughout the infection (Fig. 3B), whereas in axenic growth there was a progressive drop-off in expression after 3dpi (see Fig 2B). These results suggest that CbsR12 regulation differs in this host-cell type and that a transcriptional regulatory motif may exist outside the bounds of the *Tn327-Comp* complementation cassette, resulting in dysregulation of CbsR12 in *Tn327-Comp* compared to *Tn1832* in a THP-1 infection.

**Fig 3.**
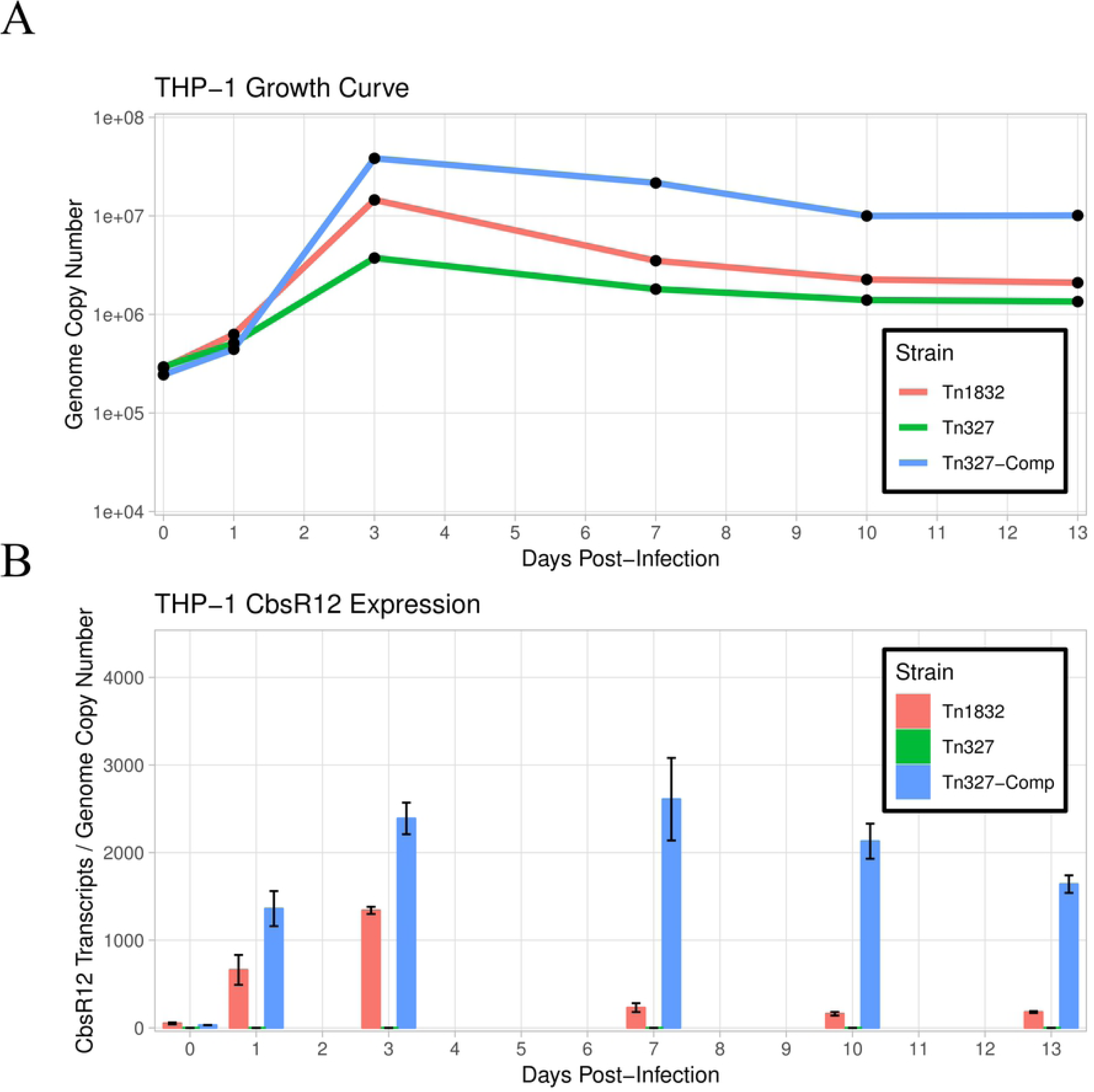
CbsR12 expression and growth effects on *C. burnetii* infecting THP-1 cells. (**A**). Growth curves for *Tn1832*, *Tn327*, and *Tn327-Comp* in THP-1 cells as determined by qPCR. The 0dpi time point refers to the inoculum. Results shown are representative of three independent experiments with consistent and indistinguishable results. (**B**). Cbsr12 expression over time for *Tn1832*, *Tn327*, and *Tn327-Comp C. burnetii* strains infecting THP-1 cells, as determined by qRT-PCR. Values represent the means ± standard error of means (SEM) of three independent determinations.

### CCV size correlates with CbsR12 expression in THP-1 infection

To more closely examine bacterial-host cell interactions, we employed immunofluorescence assays (IFAs) of *C. burnetii* infecting THP-1 cells. *C. burnetii* colonies and CCV boundaries were visualized by using anti-*Coxiella* (anti-Com1 [48]) and anti-LAMP1 antibodies at both 3dpi (late LCVs) and 7dpi (SCVs). Here, we define a *C. burnetii* colony as multiple *C. burnetii* cells inhabiting a LAMP1-decorated intracellular vacuole. LAMP1 is a host cell protein recruited to lysosomes and found on CCVs after lysosome fusion [49]. We observed a robust infection at 3dpi for *Tn1832* and *Tn327-Comp* strains, whereas the *Tn327* strain only produced a few, small CCVs with relatively unclear boundaries (Fig 4A). In contrast, the *Tn327-Comp* strain produced CCVs that were similar in size to those produced by the *Tn1832* strain, reflecting the trend observed in their respective growth curves (see Fig 3A). Quantitatively, there were significantly fewer *Tn327* colonies per THP-1 cell at 3dpi as compared to the other two strains, suggesting a possible defect in adhesion / internalization (Fig 4B). In addition, we observed significantly smaller CCVs produced by *Tn327* at 3dpi compared to the other strains (Fig 4C). However, by 7dpi there were still significantly fewer colonies in Tn327 vs. the other strains, but CCVs were of similar size in *Tn327* and *Tn1832* infections, presumably due to CCV fusogenesis. Interestingly, the *Tn327-Comp* strain formed consistently larger CCVs at 3dpi (significantly greater than *Tn1832* and *Tn327*) and 7dpi (significantly greater than *Tn327*), which meshes well with the sustained CbsR12 expression evidenced throughout the course of infection (see Fig 3B). Taken as a whole, these results suggest that CbsR12 is important for optimum growth and the establishment of CCVs early in the course of infection of THP-1 cells, and CbsR12 expression can influence CCV expansion throughout a THP-1 infection.

**Fig 4.**
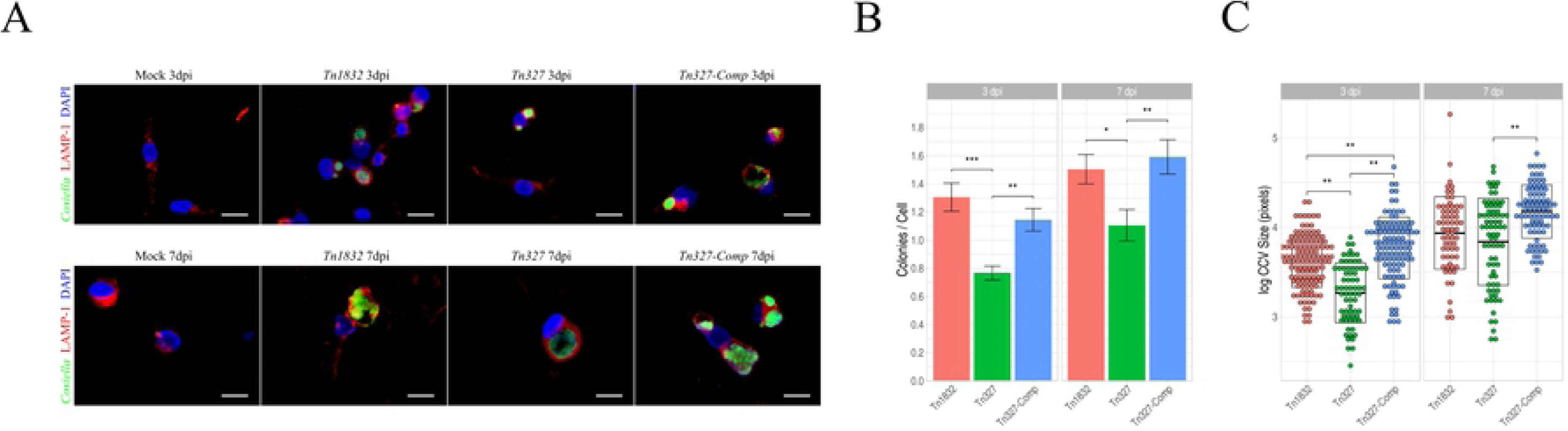
CbsR12 affects adhesion / internalization and CCV expansion in infected THP-1 cells. (**A**). Representative IFAs of *C. burnetii* transposon control strain *Tn1832*, *cbsR12* mutant strain *Tn327*, and the complemented *cbsR12* mutant strain, *Tn327-Comp,* infecting THP-1 cells at 3dpi and 7dpi. *C. burnetii* cells were probed with anti-Com1 antibodies coupled to Alexa Fluor 488 (green), CCV boundaries were labeled with anti-LAMP1 antibodies coupled to rhodamine (red), and host cell nuclei were labeled with DAPI (blue). (**B**). Number of detected *C. burnetii* colonies / THP-1 cell at 3dpi and 7dpi. Measurements were taken from 46 individual images of random fields of view from three independent experiments for each *C. burnetii* strain (* = P < 0.05, ** = P < 0.01, *** = P < 0.001, one-way ANOVA). (**C**). Sizes of individual CCVs in log_10_(pixels) for *Tn1832*, *Tn327*, and *Tn327-Comp*. Measurements were taken from 46 individual images of random fields of view spanning three different experiments for each *C. burnetii* strain. Crossbars represent the means ± standard error of means (SEM) (** = P < 0.01, one-way ANOVA). Scale bars 20 µm.

### CbsR12 binds to *carA*, *metK*, and *cvpD* transcripts *in vitro*

Although CbsR12 was identified as a CsrA-binding sRNA, nothing was known about the CsrA regulon in *C. burnetii*, making it difficult to ascribe intracellular phenotypes to regulation by CsrA. Therefore, we wanted to determine if CbsR12 can act by regulating mRNAs in *trans*. To identify potential mRNA targets of CbsR12, we first employed three *in silico* sRNA target discovery algorithms. From these search results, we omitted genes annotated as hypothetical and chose *cvpD*, *metK*, *carA*, *purH*, *rpsA*, and *dnaA* as potential targets based on conserved predictions (Table 2). To get a sense of CbsR12’s ability to bind to these potential mRNA targets, we next performed a RNA-RNA hybridization followed by EMSAs. The results clearly showed that CbsR12 bound to *carA*, *metK*, and *cvpD* transcripts *in vitro*, but did not interact with *purH*, *dnaA*, or *rpsA* mRNAs (Fig 5). We further tested CbsR12’s specificity to these transcripts by performing dose-dependent and unlabeled-chase experiments. Results of the EMSA analyses showed that CbsR12 specifically bound *carA*, *metK*, and *cvpD* transcripts in a dose-dependent manner (**S3-S5 Figs**, respectively).

**Fig 5.**
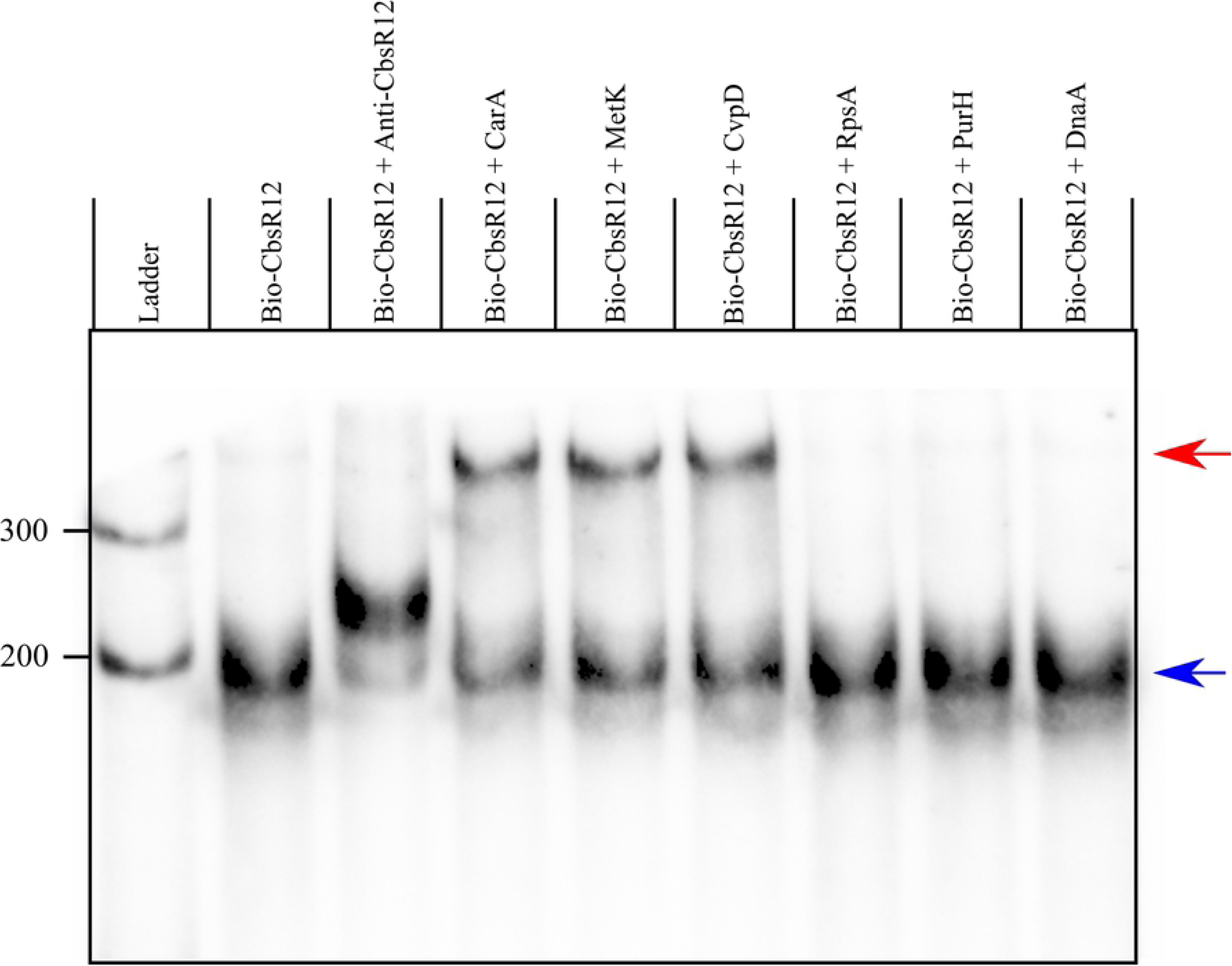
CbsR12 targets *carA*, *metK*, and *cvpD* transcripts *in vitro*. RNA-RNA EMSA showing hybridization reactions with biotin-labeled CbsR12 and *in vitro-*transcribed segments of *carA*, *metK*, cvpD, purH, *rpsA* or *dnaA*. Anti-CbsR12 represents a positive control consisting of a transcript equal in size, but antisense, to the CbsR12 transcript. Arrows indicate un-bound bio-CbsR12 (blue) and bio-CbsR12 bound to RNA targets (red).

**Table 2.**
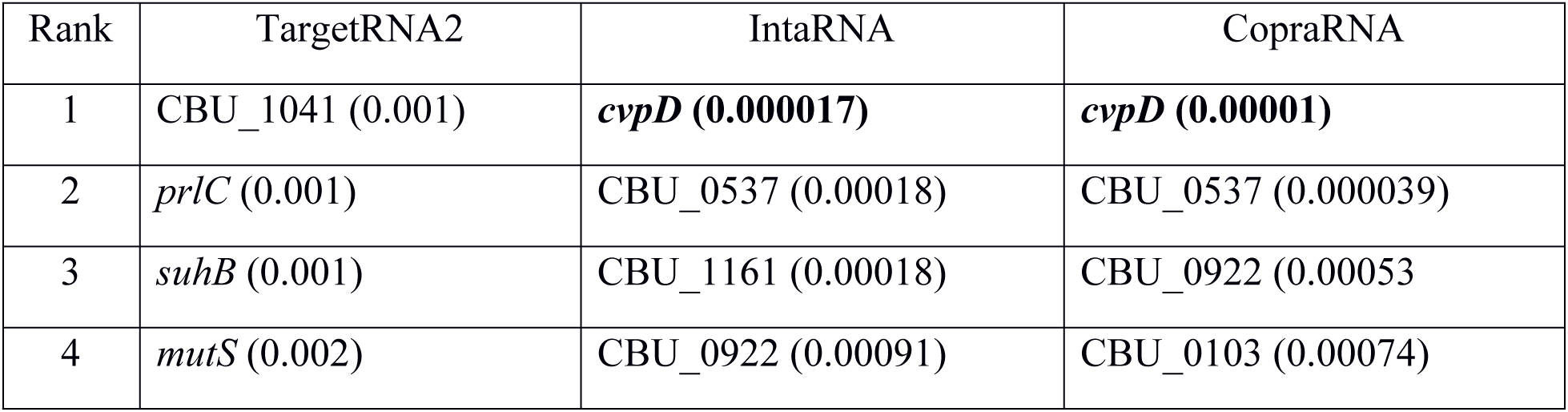
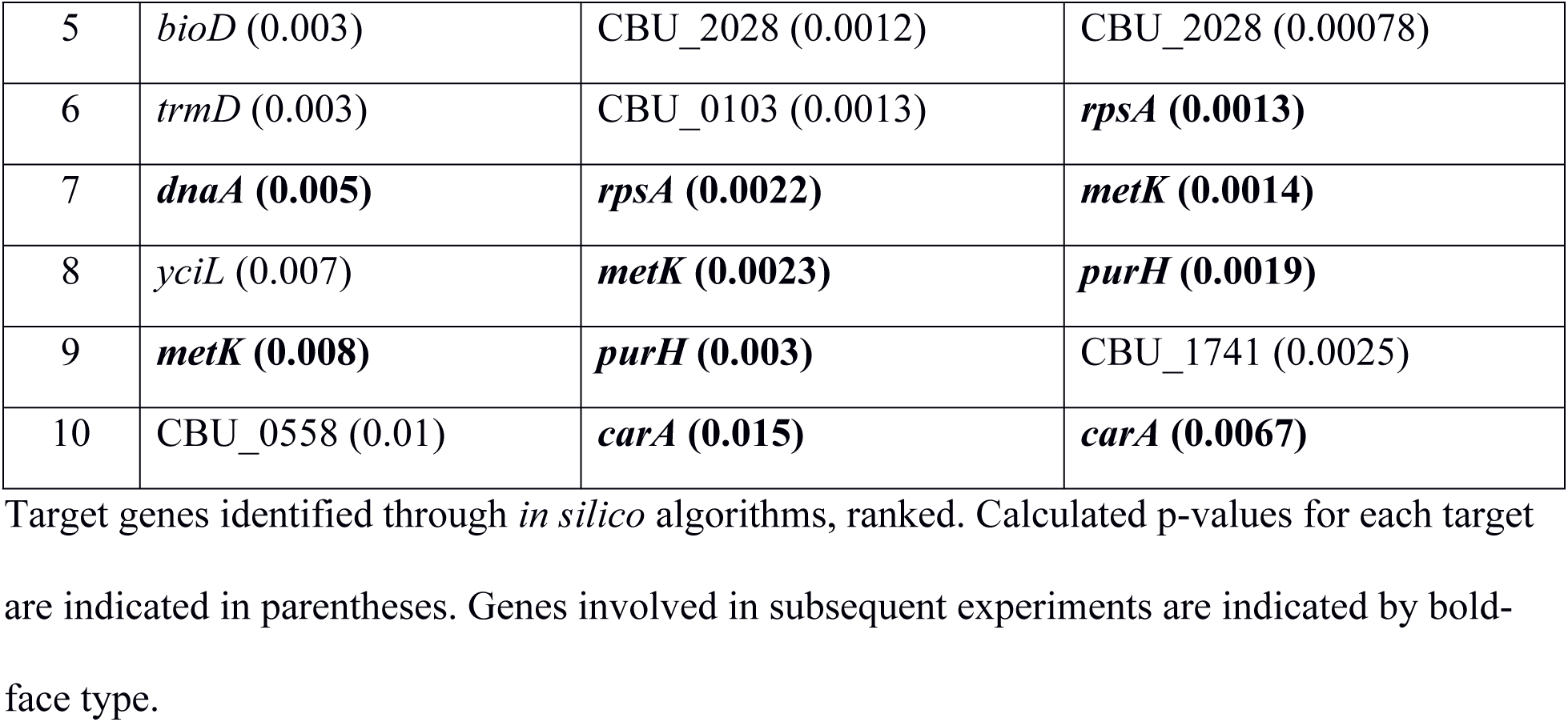
CbsR12 target prediction using various *in silico* algorithms

### CbsR12 binds to *metK*, *carA*, *cvpD*, and *ahcY* transcripts *in vivo*

To determine the CbsR12 targetome within *C. burnetii* cells, we employed a Crosslink-Seq technique previously used to detect intracellular mRNA targets of *E. coli* sRNAs [50]. For this procedure, we used *C. burnetii* LCVs grown in ACCM-2 to produce sufficient *C. burnetii* volumes to capture CbsR12 target RNAs for cDNA library preparation and RNA-Seq analysis. Hybridized RNAs in lysates from both *Tn1832* and *Tn327* strains were cross-linked, and upon RNA-Seq analysis used to identify RNAs that were enriched in *Tn1832* vs. *Tn327* strains (Fig 6). Crosslink-Seq results confirmed that CbsR12 targets *carA*, *metK*, and *cvpD* transcripts *in vivo*, as demonstrated *in vitro* (Fig 5). We also discovered an additional mRNA target, *ahcY*, coding for acylhomoserine synthase; another component of the methionine cycle. Interestingly, *ahcY* was also predicted as an mRNA target by *in silico* analyses, although the p-value was not significant (data not shown). Additionally, *ahcY* is in an operon with and downstream of *metK*. To address whether *ahcY* is actually a target of CbsR12, or if it is a result of CbsR12’s binding to a polycistronic mRNA, we used the Artemis genome browser to observe Crosslink-Seq reads aligned to the *C. burnetii* RSA439 genome. This analysis showed distinct segments of these genes to which the captured reads mapped, suggesting they are separate binding events (**S6 Fig**).

**Fig 6.**
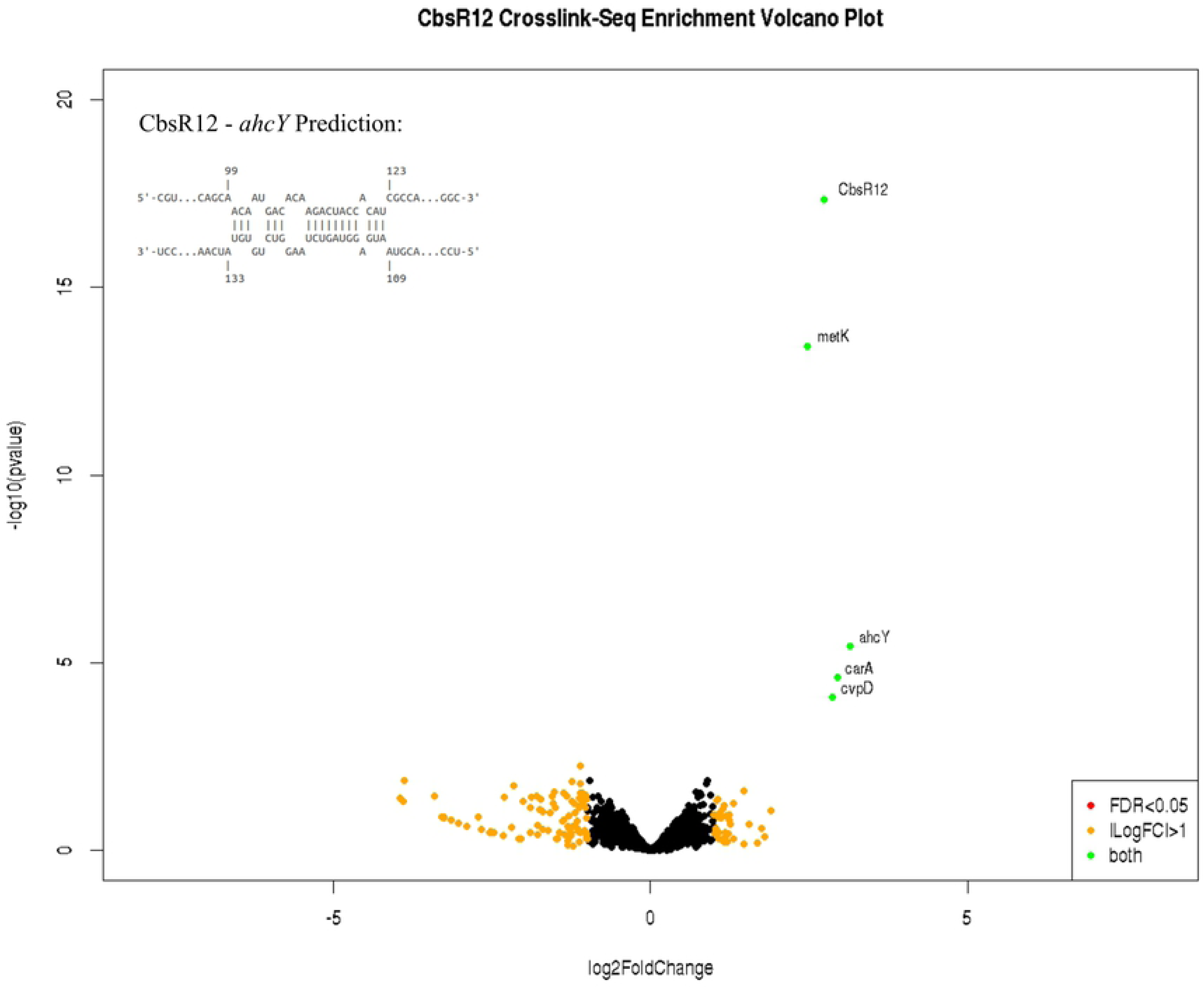
CbsR12 targets several *C. burnetii* transcripts *in vivo*, including those of *metK, carA*, and *cvpD*. Volcano plot highlighting genes that are differentially expressed between Crosslink-Seq experiments retaining CbsR12 (*Tn1832*) and lacking CbsR12 (*Tn327*). Labeled genes are indicated by a green dot and are significantly enriched in *Tn1832* versus *Tn327*, identifying them as targets of CbsR12. Black dots represent genes not differentially expressed between the conditions tested. Orange dots represent genes having a log_2_-fold change in expression > 1, but a false discovery rate (FDR) > 0.05. There were no genes indicated by red dots, although these would represent genes with a FDR < 0.05 but a log_2_-fold change in expression < 1. The potential CbsR12-binding site in the coding region of the *ahcY* transcript is inset.

### CbsR12 negatively affects the quantity of *cvpD* transcripts and regulates *carA* and *metK in vivo*

Next, we set out to determine if CbsR12 binds *carA metK*, and *cvpD* transcripts *in vivo*. First, we performed 5’ RACE on the three transcripts in total RNA extracted from *C. burnetii* LCVs infecting mammalian cells. 5’ RACE results for the *Tn1832 cvpD* gene indicated three apparent TSSs, including a TSS for the full-length transcript, questionable “TSSs” within the CbsR12-binding site, and an alternative TSS for a short transcript downstream of the CbsR12-binding site and with its own predicted promoter element (**S7A Fig**). Interestingly, putative RBSs and start codons exist downstream of TSSs for both the full-length and short transcripts. Moreover, the two start codons are in-frame with each other and the existence of putative RBSs supports the possibility that translation occurs from both elements. The questionable “TSSs” within the CbsR12-binding region likely result from CbsR12-mediated RNase III degradation of *cvpD* mRNA, because 5’ RACE results for *Tn327*, lacking Cbsr12, did not produce TSSs in this region (data not shown). We predict that CbsR12 down-regulates expression of full-length CvpD since the CbsR12-binding site occurs in the coding region. However, CbsR12 would predictably have no effect on expression of the putative truncated CvpD, as the CbsR12 binding site occurs upstream of the alternative TSS (**S7A Fig**).

To determine if CbsR12 binds to and causes degradation of full-length *cvpD* transcripts, we also performed qRT-PCR on *Tn1832*, *Tn327*, and *Tn327-*Comp LCVs obtained from infected THP-1 cells. These results clearly showed that the absence of CbsR12 in strain *Tn327* leads to a significant increase in full-length *cvpD* transcripts in LCVs (3dpi) (**S7B Fig**). At 7dpi, *Tn1832* and *Tn327* levels were not significantly different, presumably due to reduced CbsR12 expression in *Tn1832* SCVs (see Fig 3B). However, *Tn327-Comp cvpD* expression was significantly lower, most likely due to the maintained expression of CbsR12 in *Tn327-Comp* SCVs (see Fig 3B). Whether or not two different forms of CvpD protein are produced from the *cvpD* gene is unknown, although it appears that CbsR12 may negatively regulate the full-length *cvpD* transcript *in vivo*.

In order to determine if CbsR12 binds to *carA* and *metK* transcripts *in vivo*, we devised a novel reporter assay in *E. coli*. 5’ RACE results for both *Tn1832* and *Tn327* strains revealed that *carA* has two potential TSSs, a finding that is consistent with transcription of *E. coli carA* [42]. Based on the position of the TSSs, CbsR12 could only regulate the longer, full-length *carA* transcript and not the shorter mRNA, whose transcription starts immediately upstream of the ribosome-binding site (RBS) and downstream of the CbsR12-binding site (Fig 7A). From these results, we hypothesized that CbsR12 binds to the 5’ untranslated region (UTR) of *carA* and upregulates translation by relieving the secondary structure that occludes the predicted RBS (Figs 7A, 7B). Results of the *E. coli* reporter assay confirmed our hypothesis, because translation of luciferase enzyme from a *carA5’UTR*-*luc* fusion was significantly upregulated in the presence of CbsR12 relative to a strain lacking the sRNA (Fig 7C).

**Fig 7.**
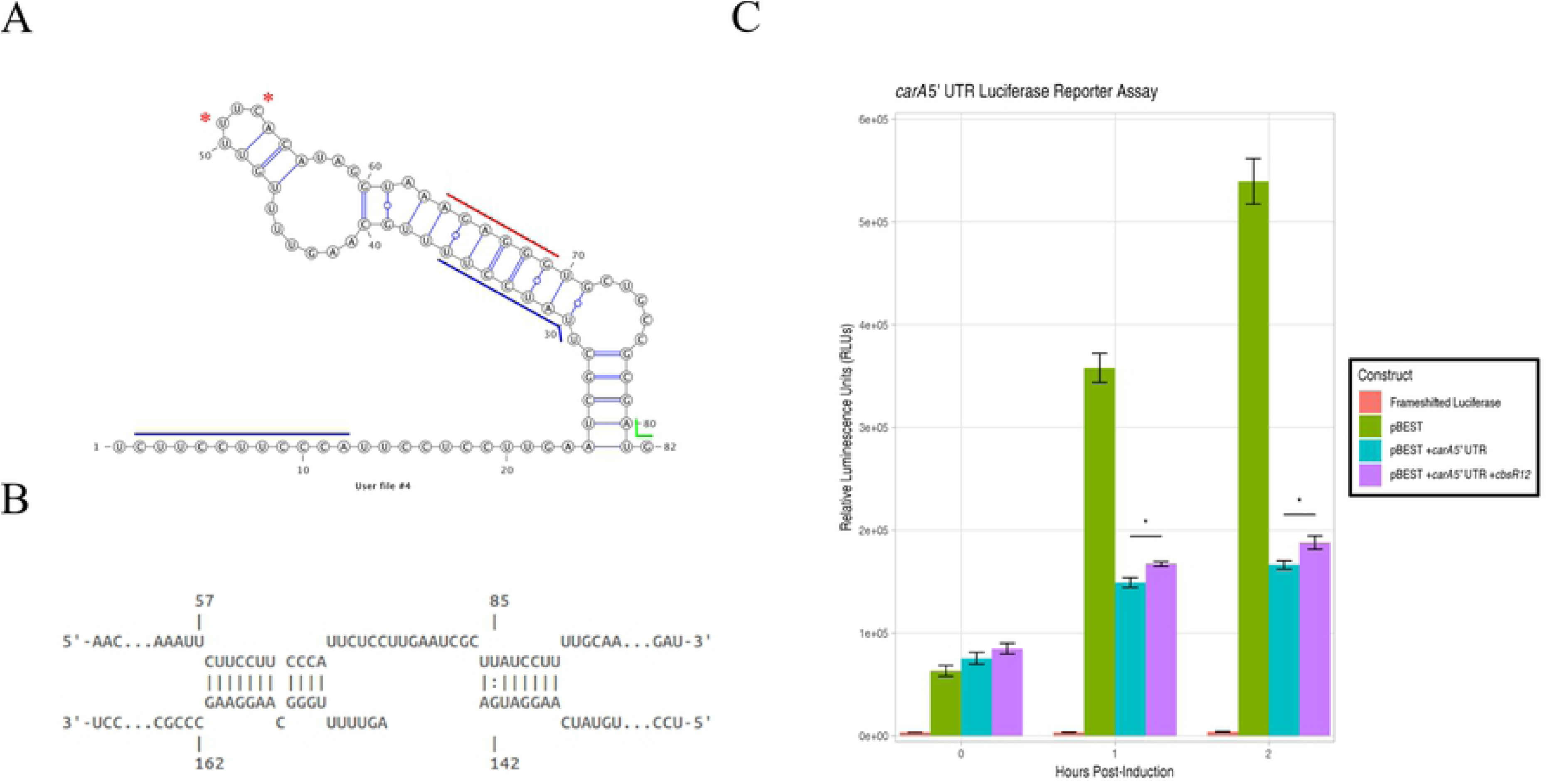
CbsR12 targets and upregulates expression of a *carA-*luciferase translational fusion construct *in vivo*. (**A**). Secondary structure of the *carA* 5’ UTR as predicted by mFold. Red asterisks indicate the TSSs for the shorter transcripts as determined by 5’ RACE. (Nucleotide 1 was determined to be the TSS for the full-length transcript by 5’ RACE). Colored lines represent the start codon (green), predicted RBS (red), and determined CbsR12-binding sites (blue). (**B**). Representation of CbsR12 binding to the *carA* transcript as determined by IntaRNA, with respective base numbers indicated. The top strand in the model represents the *carA* sequence, while the bottom strand represents the complementary CbsR12 sequence. (**C**). *carA*-*luc* reporter assay indicating relative luminescence units produced by pBESTluc constructs with: 1) no luciferase expression (Frameshifted Luciferase), 2) pBESTluc vector (pBEST), 3) pBESTluc with the *carA* 5’ UTR upstream of *luc* but lacking *cbsR12* (pBEST + *carA* 5’ UTR), and 4) *carA* 5’ UTR upstream of *luc* plus the *cbsR12* gene driven by a P*tac* promoter (pBEST + *carA* 5’ UTR + *cbsR12*) (* = P < 0.05, student’s t-test).

In contrast, CbsR12 was predicted to downregulate MetK translation by binding to the coding region of the transcript, immediately downstream of its start codon (Fig 8A). As is often the case with this type of sRNA-mediated regulation, RNase III would likely be recruited and the *metK* transcript cleaved, resulting in downregulation of the encoded protein product. Unexpectedly, 5’ RACE analyses of *metK* mRNA also identified apparent alternative “TSSs” within the CbsR12-binding region, suggesting that the truncated mRNAs resulted from CbsR12-mediated RNase III processing (Figs 8A, 8B). Indeed, 5’ RACE analysis of RNA from strain *Tn327* infecting THP-1 cells did not detect the “TSSs”, suggesting they are a product of RNase III processing (data not shown). Results of the reporter assays in *E. coli* confirmed our hypothesis, as the presence of CbsR12 significantly down-regulated translation of luciferase enzyme from the *metK*-*luc* fusion construct compared to a strain lacking the sRNA (Fig 8C).

**Fig 8.**
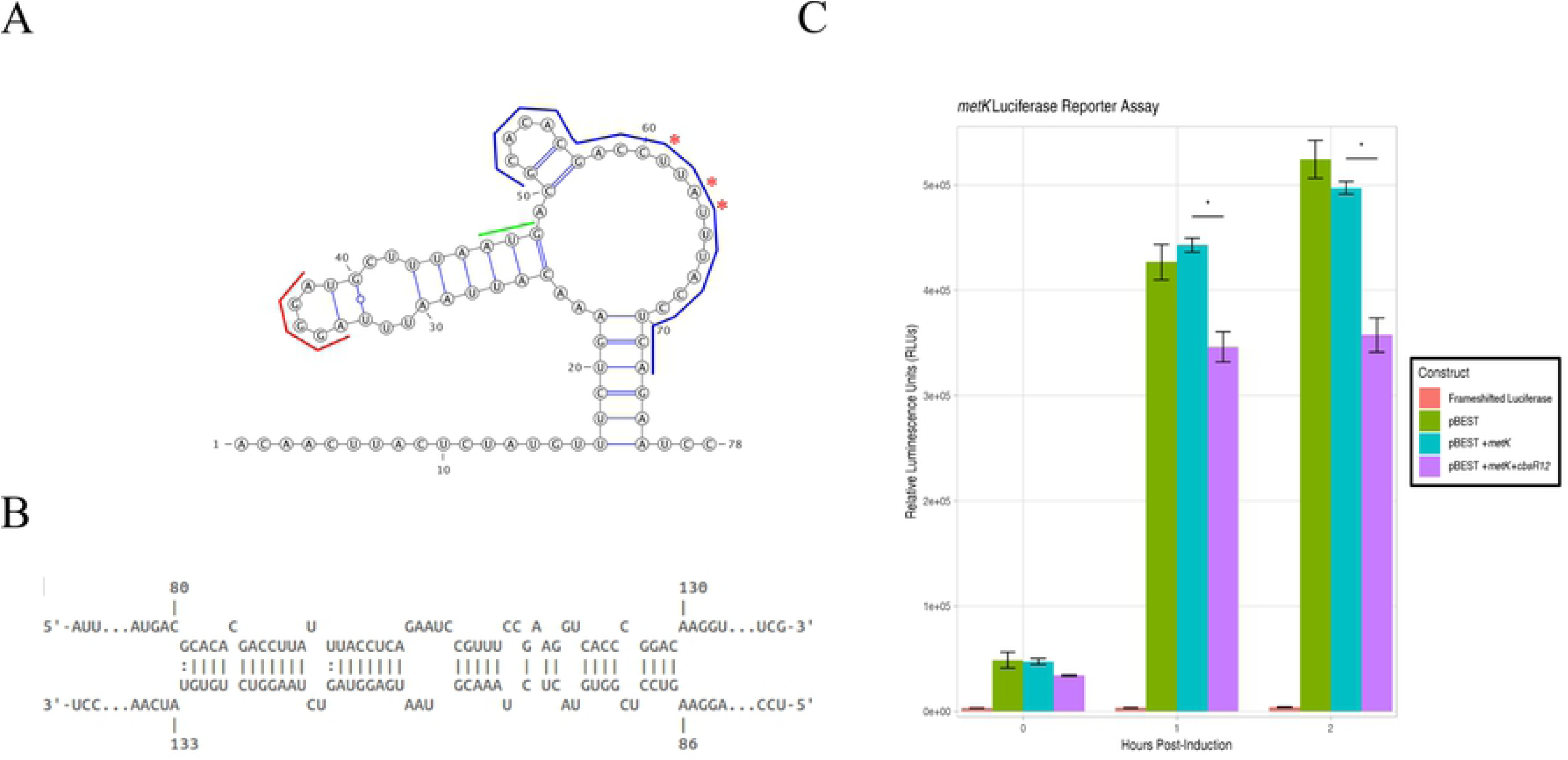
CbsR12 targets and downregulates expression of a *metK*-luciferase translational fusion construct *in vivo*. (**A**). Secondary structure of the *metK* 5’ UTR and initial coding sequence as predicted by mFold. Red asterisks indicate apparent alternative “TSSs” determined by 5’ RACE. (Nucleotide 1 was determined to be the TSS for the full-length transcript by 5’ RACE). Colored lines represent the start codon (green), a predicted RBS (red), and the determined CbsR12-binding site (blue). (**B**). Representation of CbsR12 binding to the *metK* transcript as determined by IntaRNA with base numbers indicated. The top strand in the model represents the *metK* sequence, while the bottom strand represents the complementary CbsR12 sequence. (**C**). *metK*-*luc* reporter assay indicating relative luminescence units produced by pBESTluc constructs with: 1) no luciferase expression (Frameshifted Luciferase), 2) pBESTluc vector (pBEST), 3) pBESTluc with the CbsR12 binding site cloned in frame into *luc* but lacking the *cbsR12* gene (pBEST + *metK*) and 4) pBESTluc with the CbsR12 binding site cloned in frame into *luc* plus the *cbsR12* gene driven by a P*tac* promoter (pBEST + *metK* + *cbsR12*) (* = P < 0.05, student’s t-test).

Although we determined that CbsR12 targets *carA* and *metK* transcripts *in vitro* and *in vivo*, we were curious whether the absence of CbsR12 would also result in differential amounts of CarA and MetK proteins *in vivo* in *C. burnetii*. To this end, we performed Western blots with monospecific polyclonal antibody probes generated against recombinant *C. burnetii* CarA and MetK. As predicted, when proteins from whole-cell lysates of *Tn1832*, *Tn327*, and *Tn327-Comp* strains were compared, we found that CarA was expressed in the control (*Tn1832*) and complemented (*Tn327-Comp*) strains at comparable levels, but was undetectable in protein profiles of the *cbsR12* mutant (strain *Tn327*) (Fig 9A). In sharp contrast, MetK was highly expressed in the *cbsR12* mutant (strain *Tn327*) but expressed at relatively lower and comparable levels in the *Tn1832* and *Tn327-Comp* strains (Fig 9B).

**Fig 9.**
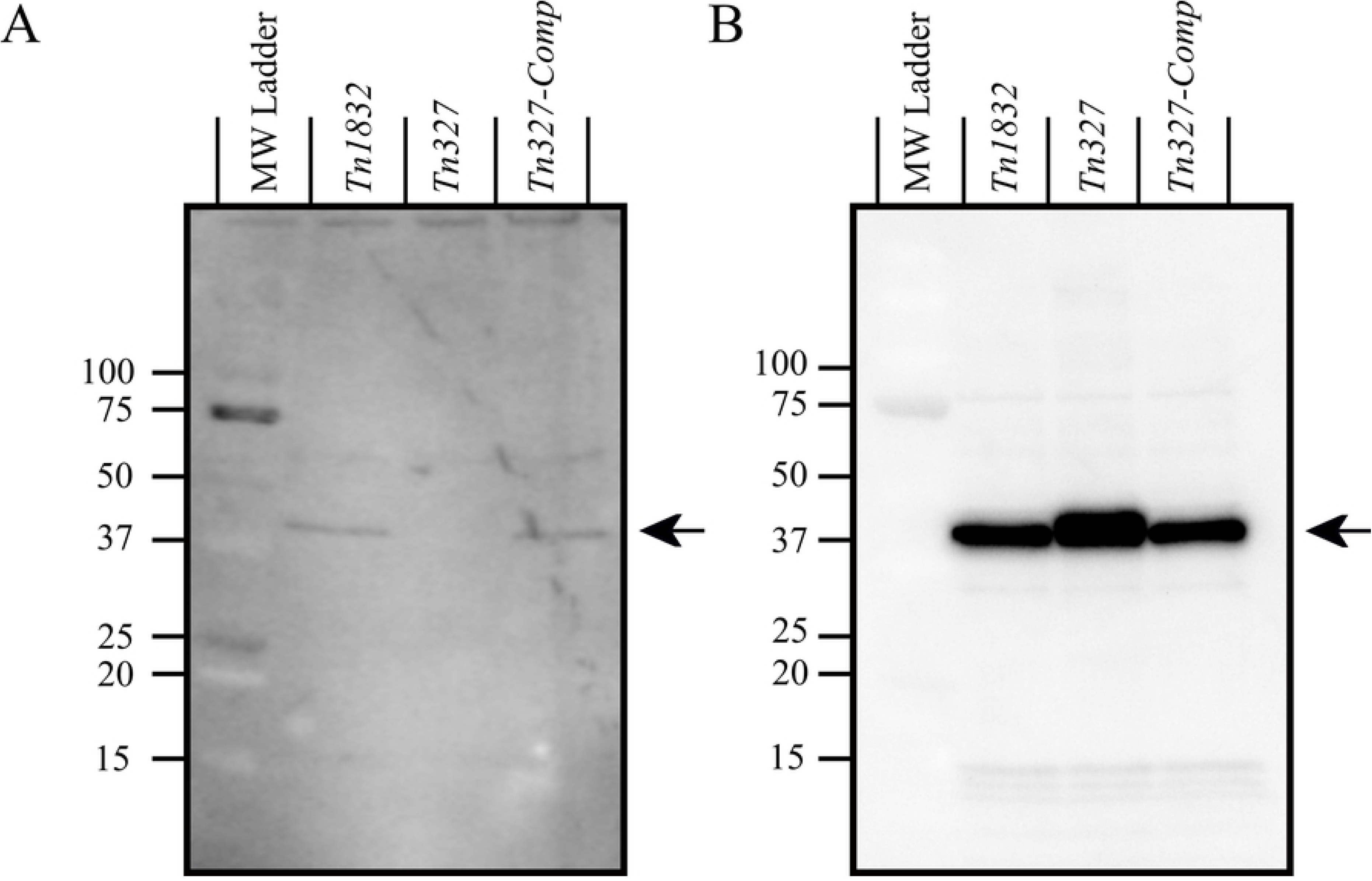
CarA and MetK proteins are differentially expressed between *Tn1832*, *Tn327*, and *Tn327-Comp*. (**A**). Proteins (30 µg total) from *Tn1832*, *Tn327*, and *Tn327-Comp* LCVs (96h for *Tn1832* and *Tn1832-Comp* and 144h for *Tn327*) grown in ACCM-2 were resolved on a 10-20% Tris-glycine SDS-PAGE gel, blotted, probed with rabbit anti-CarA antibodies, and detected with chemiluminescence. Black arrow indicates *C. burnetii* CarA. (**B**). Proteins (60 µg total) from *Tn1832*, *Tn327*, and *Tn327-Comp* LCVs (96h for *Tn1832* and *Tn1832-Comp* and 144h for *Tn327*) grown in ACCM-2 were resolved on a 10-20% Tris-glycine SDS-PAGE gels, blotted, probed with rabbit anti-MetK antibodies, and detected with chemiluminescence. The black arrow indicates *C. burnetii* MetK.

## Discussion

Considerable effort has gone into elucidating *C. burnetii* virulence factors in recent years, with much of that activity revolving around the discovery and characterization of the repertoire of T4BSS effectors that are necessary for adherence, invasion, and establishing the intracellular niche [reviewed in [51]]. Here, we present a different approach to identifying *C. burnetii* factors that potentiate a successful infection. Namely, we have focused our efforts on elucidating the roles of sRNAs in *C. burnetii*. In the present study, we show that CbsR12, a multifunctional sRNA that binds to mRNAs and to the regulatory protein CsrA-2, is important for proper *Coxiella* replication and CCV expansion during infection of human macrophage-like phagocytic cells.

We initially became interested in CbsR12 based on its remarkably high expression in LCVs and SCVs obtained from infected Vero cells relative to *C. burnetii* grown axenically in ACCM-2 (SRP041556). Induction of CbsR12 expression in *in vivo* vs. *in vitro* conditions led us to label CbsR12 as “infection-specific”. Indeed, in a *C. burnetii* infection of Vero cells, CbsR12 represents the dominant non-tRNA/rRNA/tmRNA transcript in both LCVs and SCVs. This notion is corroborated by RNA-Seq data obtained from *C. burnetii* LCVs infecting THP-1 cells (see Table 1). Collectively, these results indicate that CbsR12 is upregulated under intracellular conditions; most likely by means of a transcriptional regulator. The CbsR12 DNA sequence itself is conserved among all *C. burnetii* strains sequenced to date, underscoring the potential for an important regulatory role, but from an evolutionary viewpoint. Interestingly, though, the CbsR12 sequence is missing or degenerate in *Coxiella*-like endosymbionts [52]. This observation suggests that the sRNA is important for a mammalian infection but is dispensable in *Coxiella*-like endosymbionts that reside in arthropods.

*C. burnetii* has several potential transcription factors that are known to upregulate expression in other bacteria, including Integration Host Factor (IHF) [53], response regulator PmrA [54], and transcription factor DksA [55]. However, little is known about the roles for IHF and DksA in *C. burnetii*. The consensus IHF-binding site occurs in a *C. burnetii* selfish genetic element, suggesting that this binding sequence is maintained in the pathogen [31]. Similarly, a role for DksA in *C. burnetii*’s stringent response has been suggested, although no correlations could be drawn [56]. Only PmrA has been well studied, to date, among putative transcription factors of *C. burnetii* [19]. It is interesting to note that the *cbsR12* gene contains a close approximation to a PmrA-binding site (consensus sequence with less conserved nucleotides in lowercase: cTTAA-N_2_-TT-N_2_-cTTAA) [57] immediately upstream of its predicted −10 promoter element (CbsR12 sequence: gTTTA-N_2_-TT-N_1_-gTTAA). However, the presence of this sequence does not explain the prolonged CbsR12 expression observed during a THP-1 infection by *Tn327-Comp* since the putative PmrA-binding sequence is present in the *cbsR12* gene cassette that was cloned into *Tn327-Comp*.

The apparent down-regulation of CbsR12 after LCV-to-SCV morphogenesis occurs can possibly be explained by the smaller, CbsR12 RNA fragment that appears in roughly equal abundance to that of the full-length transcript in SCVs [29]. Results of 5’ RACE analyses in this study suggest that the apparent “TSSs” of CbsR12 occur in a double-stranded RNA stem structure, and that RNase III processing may actually act to reduce the levels of full-length CbsR12 (see **S1 Fig**). We hypothesize that RNase III-mediated decay is involved in the downregulation of CbsR12 as infection proceeds and the LCV-to-SCV transition occurs.

In this study, we have highlighted the effects of CbsR12 expression levels during a *C. burnetii* infection of the THP-1 cell line. For example, CbsR12 seems to be important for adhesion / internalization by phagocytic THP-1 cells (see Fig 4B). *C. burnetii* adherence to THP-1 cells is mediated by αVβ3 integrins, so it is conceivable that CbsR12 expression affects adherence to and invasion of THP-1s, perhaps indirectly via CsrA sequestration [58]. Additionally, the serendipitous dysregulation of CbsR12 in the *Tn327-Comp* strain allowed us to observe the effects of CbsR12 expression on growth rate and CCV sizes during infection. For example, in THP-1 cells, we saw a correlation between CbsR12 expression and growth rate from 1dpi to 3dpi. Furthermore, we found that maintenance of CbsR12 expression as the *C. burnetii Tn327-Comp* infection proceeded led to significantly larger CCVs at 7dpi. The apparent dysregulation of CbsR12 in the *Tn327*-Comp strain during infection of THP-1 cells is possibly due to the genomic context of the *Tn327*-Comp transposon-mediated insertion of *cbsR12*. We confirmed that the *cbsR12* cassette inserted into an intergenic region between CBU_1788 and CBU_1789 (RSA493: accession number NC_002971.4), and copy number qPCR analysis showed that the cassette only inserted once (data not shown). Although this *cbsR12* cassette contained the *cbsR12* gene and ∼100-bp flanking regions, it is possible that a regulator exists >100bp upstream of the *cbsR12* gene, and that this regulatory element is missing in the *Tn327*-Comp strain.

We also sought to elucidate the role for CbsR12 during infection. In doing so, we have shown that CbsR12 binds to several mRNA transcripts in *trans*. For example, we found that CbsR12 binds *carA* transcripts (see Fig 5, 7) and upregulates expression of *C. burnetii* CarA (carbamoyl phosphate synthetase A) (see Fig 9A), which is the first committed step in pyrimidine and arginine biosynthesis [42]. Pyrimidine metabolism in *C. burnetii* presumably requires CarAB to catalyze the conversion of L-glutamine into carbamoyl phosphate and glutamate, since it is unable to shunt this process through the arginine dihydrolase pathway; *C. burnetii* apparently lacks the necessary enzymes [59]. CbsR12-mediated upregulation of CarA in LCVs would result in increased production of pyrimidines that the pathogen requires for robust intracellular growth.

In *E. coli*, *carA* expression is tightly controlled by a series of transcriptional regulators and the presence of two distinct promoters that are regulated by feedback from arginine and pyrimidines [42]. 5’ RACE analysis showed two distinct TSSs for *C. burnetii carA* mRNA, with the full-length transcript containing two CbsR12-binding sites and a shorter putative transcript lacking both of them (see Fig 7A). We do not believe that the shorter, alternative TSS is due to RNase III-mediated degradation resulting from CbsR12 binding because this alternative TSS remained in 5’ RACE analysis of the *Tn327* strain (data not shown). We do not know the conditions under which the shorter transcript is expressed, but it may involve feedback from arginine/pyrimidine in accordance with *carA* regulation in *E. coli*.

We also showed that CbsR12 binds to *metK* transcripts (see Fig 5, 8) and downregulates expression of *C. burnetii* MetK protein (see Fig 9B). MetK is a key component of the methionine cycle and serves to synthesize SAM, the major methyl donor in bacterial cells, from a methionine substrate. The methionine cycle is a series of reactions catalyzing methionine to SAM via MetK, SAM to S-adenosylhomocysteine (SAH) via various methylases, SAH to homocysteine via AhcY, and homocysteine to methionine via MetH/MetE. Cells produce homocysteine as an input molecule through a series of reactions involving activated homoserines [reviewed in [60]]. *C. burnetii* is considered to be a semi-auxotroph for methionine, since it can potentially grow without methionine in axenic media, albeit at a slower growth rate [61]. Interestingly, *C. burnetii* seems to lack several components of the methionine synthesis pathway, most notably the ability to produce activated homoserines. Most bacteria activate homoserine through an O-succinyl group catalyzed by MetA or an O-acetyl group catalyzed by MetX [reviewed in [60]]. *C. burnetii* apparently lacks genes coding for these proteins. An ABC methionine transporter has been hypothesized [61] but not verified in *Coxiella*. If this is indeed a functional transporter, CbsR12’s negative regulation of *metK* transcripts makes sense in the context of the sRNA’s high expression in LCVs, because any amount of scavenged methionine would be critical to growth. Shifting the equilibrium from SAM synthesis to methionine retention would be necessary, at least initially, as *C. burnetii* rapidly produces proteins to expand its intracellular niche.

SAM is also a crucial molecule contributing to most bacterial processes [62]. Specifically, as the major methyl donor, it is necessary for regulation of numerous enzymes and has been implicated as a major contributor to virulence [63]. Some bacteria lack a functional *metK* gene and instead transport SAM directly [64]. There are many uncharacterized transporters encoded in the *C. burnetii* genome, so it is conceivable that a SAM transporter is present [65]. This would allow for SAM scavenging even when MetK production is downregulated by CbsR12. Furthermore, assuming that SAM transport occurs, *C. burnetii* could synthesize methionine without having to scavenge it, since methionine can be synthesized from SAM without activated homoserine via the methionine cycle.

Crosslink-Seq (see Fig 6) also determined that CbsR12 actively targets *ahcY* transcripts coding for adenosylhomocysteinase, another component of the methionine cycle. AhcY catalyzes SAH into homocysteine and adenosine. Based on the location of the CbsR12-binding site in the coding region of the *ahcY* transcript (Fig 6 **inset**), we predict that CbsR12 negatively regulates AhcY expression. The reason for this negative regulation is currently unknown, although it could be involved in a mechanism to suppress adenosine and/or homocysteine accumulation in LCVs.

*cvpD* was also determined to be a CbsR12 target through Crosslink-Seq (Fig 6), and this was confirmed by RNA-RNA hybridization / EMSA and qRT-PCR analyses (see Fig 5**, S7B Fig**). In this study, we found that CbsR12 was necessary for CCV expansion in early stages of a THP-1 infection. The mechanism for this is unclear, although it may revolve around regulation of *cvpD*, which was shown to be required for *C. burnetii*’s intracellular replication and CCV expansion in infected THP-1 and HeLa cells [11]. CbsR12 is predicted to target the coding region of the *cvpD* transcript and thus negatively regulate translation. However, in the context of CbsR12’s expression pattern, this is an unclear connection, as one would expect upregulation of CvpD synthesis at a time when CbsR12 is highly expressed in LCVs. However 5’ RACE analysis of *cvpD* in *Tn1832* and *Tn327* provides a potential explanation, as there appears to be an alternative *cvpD* promoter downstream of the CbsR12-binding site that also possesses a putative RBS and start codon (see **S7A Fig**). From these results, we hypothesize that there are two gene isoforms of *cvpD* that are transcribed and differentially expressed based on the *C. burnetii* morphotype. Due to high expression of CbsR12 in LCVs, the longer *cvpD* transcript isoform would be downregulated by RNase III-mediated decay. As expression of CbsR12 decreases as the infection proceeds, the longer transcript isoform would be able to accumulate. qRT-PCR data support this explanation, since a lack of CbsR12 in the *Tn327* mutant increased the amount of transcripts corresponding to the long *cvpD* isoform (see **S7B Fig**). This hypothesis could be confirmed through Western blot analysis if the two putative CvpD products could be identified and distinguished.

A recent study on the 2007-2010 Dutch outbreak of *C. burnetii* yielded several newly annotated *C. burnetii* genomes specific to that epidemic [66]. Curiously, 7 of the 13 strains analyzed contained a frameshift deletion in the *cvpD* gene, leading to premature stop codons [66]. Among these, strains 18430 (NZ_CP014557.1), 14160-001 (NZ_CP014551.1), 701CbB1 (NZ_CP014553.1), and 2574 (NZ_CP014555.1) had single-base deletions that only affected the long *cvpD* isoform. These strains were all isolated from aborted placentas of ruminants and cattle in the Netherlands and France [66]. Additional related strains, but not isolated from the Dutch outbreak, include the Heizberg (NZ_CP014561.1), Henzerling (NZ_CP014559.1), and RSA 331 (NC_010117.1). These strains, which were isolated from patients with acute Q fever in northern Italy and Greece in the mid 1900’s, harbor 4-bp frameshift deletions towards the middle of the *cvpD* coding region, affecting both long and short *cvpD* isoforms and introducing premature stop codons [66, 67]. Apparently, CvpD was dispensable for virulence in the latter strains, while the recent Dutch outbreak strains harbored either intact *cvpD* genes, or *cvpD* genes with a 1-bp frameshift deletion only affecting the longer gene isoform. From these observations, it is tempting to speculate on the utility of *cvpD* in *C. burnetii*. A *cvpD* deletion in *C. burnetii* RSA439 produced an intracellular phenotype very similar to the *Tn327* phenotype [11], although in the aforementioned strains of *C. burnetii* in which *cvpD* is pseudogenized, CbsR12 regulation of the gene is apparently dispensable. Granted, there are many genotypic differences between RSA439 and the Dutch isolates [66], and some compensatory mechanism(s) may exist for the absence of *cvpD*. Alternatively, *cvpD* may be necessary during infection of human cell lines and dispensable in host-animal infections. Regardless, the role and regulation of the CvpD effector requires further study.

This study also highlights the existence of CsrA-binding sites in the secondary structure of CbsR12 (see **S1A Fig**). We showed that CbsR12 binds rCsrA-2, but not rCsrA-1, in a dose-dependent manner, *in vitro.* It is interesting that CsrA-1 does not bind to the consensus motifs present in CbsR12, and the reason for this is unclear, especially since both CsrA-1 and CsrA-2 maintain the critical L4 and R44 RNA-binding residues, although CsrA-1 has these residues at L4 and R46 [68]. There are several examples of pathogens harboring multiple copies of CsrA [69, 70]. For example, RsmF of *P. aeruginosa* is a homolog of RsmA (CsrA) and functions by binding a subset of mRNAs that RsmA also binds [70]. However, a *rsmF* mutant did not display a phenotype during infection [70]. Similarly, In *C. burnetii*, a transposon-mediated *csrA-1* mutant was shown to have no intracellular phenotype [15]. It is also conceivable that CsrA-1 has diverged during *C. burnetii*’s adaptation to an intracellular lifestyle and is no longer functional. It is also possible that CsrA-1 binds to a non-canonical motif that is not present in CbsR12, although such CsrA homologs have not yet been described, to our knowledge. Regardless, it is necessary to examine the CsrA-1 and CsrA-2 regulons in order to determine their respective roles during infection.

In *L. pneumophila*, a close relative of *C. burnetii*, successful infection depends on a LetAS-RsmYZ-CsrA regulatory cascade. LetAS is a two-component system (TCS) that regulates expression of two sRNAs, RsmY and RsmZ, which in turn act as RNA “sponges”, to soak up the CsrA RNA-binding global regulatory protein and modulate its activity [41]. We determined that CbsR12 binds *C. burnetii* rCsrA-2 *in vitro*, and requires only four CsrA-binding sites, in a fashion similar to the RsmY/Z CsrA-binding sRNAs of *L. pneumophila*. Interestingly, *L. pneumophila* RsmY/Z was implicated in the formation of cell aggregates and biofilms when the sRNAs were ectopically overexpressed in *E. coli*, mimicking the effects of *E. coli*’s own CsrA-binding sRNAs [39]. Likewise, when we overexpressed CbsR12 in *E. coli* during reporter assays (Figs 7, 8), we observed a similar autoaggregative phenotype (**S8 Fig**).

Since it is likely that CbsR12 serves as a *C. burnetii* RsmY/Z homolog, it is worth noting that we have identified a second sRNA, ***<u>C</u>****oxiella **<u>b</u>**urnetii* **<u>s</u>**mall **<u>R</u>**NA **1** (CbsR1), that harbors 5 putative CsrA-binding sites with the ANGGA motif (data not shown) [29]. Similar to Cbsr12, CbsR1 (another sRNA) is also highly expressed in *C. burnetii* LCVs infecting Vero cells (see Table 1). Furthermore, *cbsR1* harbors a putative LetA-binding site similar to that of *L. pneumophila* RsmY (data not shown) [39]. Together, these observations suggest that CbsR1 and CbsR12 may represent orthologs of RsmYZ, although further exploration of CbsR1 and its cooperativity with CbsR12 is required. If CbsR1 does indeed serve as a CsrA-binding sRNA, its potential, redundant regulatory role may help to explain why *Tn327* CCVs expanded to wild-type sizes as the infection progressed (see Fig 4C).

CbsR12’s high level of expression during infection likely facilitates and regulates the many functions we have described. We predict that expression of *cbsR12* is itself regulated by an unidentified TCS in a fashion similar to the *L. pneumophila* LetAS TCS regulation of RsmYZ sRNAs [41]. In fact, the LetAS TCS of *L. pneumophila* is not unlike the GacAS TCS involved in RsmYZ-CsrA cascades in other bacteria [71]. *C. burnetii* codes for three different GacA response regulators that could bind upstream elements of the *cbsR12* gene and regulate expression [65]. This upstream regulator may, in turn, help to explain the dysregulation of expression seen in *Tn327-Comp* during infection of THP-1 cells (see Fig 2B) that is not apparent during axenic growth (see Fig 1B). Alternatively, CbsR12 expression could be upregulated by PmrA, and some other regulator may be involved in its down-regulation, in conjunction with RNase III-mediated decay. Together, these would aid in suppression of CbsR12 as the LCV-to-SCVs transition occurs, effectively freeing sequestered CsrA-2 to regulate the fate of its target transcripts.

Free CbsR12 not bound to CsrA-2 presumably acts in *trans* to facilitate efficient replication through translational up-regulation of CarA and down-regulation of MetK, and perhaps by enabling expansion of the CCV by means of *cvpD* transcript regulation. It is worth noting that the CsrA-binding sites of CbsR12 do not overlap with *metK* and *cvpD* binding sites. Hence, it is feasible, that CbsR12 may still regulate *metK* and *cvpD* in *trans* while bound to CsrA-2; in fact, a chaperone-like function such as this has recently been ascribed to CsrA [72].

CbsR12 is one of few identified *trans*-acting sRNAs that also binds CsrA [reviewed in [73]]. For example, McaS in *E. coli* acts in *trans* to regulate flagellar biosynthesis and biofilm formation [74]; however, some of these functions could be due to McaS-mediated sequestration of CsrA rather than via in-*trans* targeting [75]. Similarly, CbsR12’s role in regulating *C. burnetii* replication and CCV expansion could be due to a combination of in-*trans* mRNA (*metK*, *carA*, *cpvD*) targeting and regulation of CsrA-2 function. Currently, the CsrA regulon in *C. burnetii* is completely unknown, so it is not possible to ascribe these phenotypes to a CbsR12-CsrA regulatory cascade. Our lab is currently working to elucidate the CsrA-1/CsrA-2 regulons, along with regulation of the putative CbsR12-CsrA-2 cascade of *C. burnetii* and the nature of CsrA-CbsR1 binding.

## Materials and Methods

### Bacterial strains, cell lines and growth conditions

Strains, primers, and plasmids used in this study are listed in **S9 Fig**. *E. coli* was grown in lysogeny broth (LB) supplemented with ampicillin (100 µg/mL) or kanamycin (50 µg/mL), as needed. When necessary, overnight *E. coli* cultures were expanded to 100 mL LB, grown for two hours, then supplemented with 1 mM IPTG for induction. *C. burnetii* Nine Mile phase II (strain RSA439, clone 4), *Tn1832*, *Tn327*, and *Tn327-Comp* were grown in ACCM-2 medium [30] supplemented with ampicillin (5 µg/mL) or kanamycin (350 µg/mL) at 5% CO_2_, 2.5% O_2_, 92.5% N_2_ 37°C, and 100% humidity with continuous shaking at 75 RPM [30]. SCVs collected from Vero cells were used for all *C. burnetii* infections and growth curve experiments. Briefly, *C. burnetii* was used to infect Vero cell monolayers for seven days at 5% CO_2_ and 37°C for seven days, after which the cultures were removed to room temperature and the flask lids tightened and covered for two additional weeks [76]. Following this, SCVs were harvested with digitonin, as previously described [77].

African green monkey kidney (Vero) epithelia (CCL-81; American Type Culture Collection; ATCC) and human monocytic leukemia (THP-1) cells (TIB-202; ATCC) cell lines were maintained in RPMI medium (Gibco) supplemented with 10% fetal bovine serum (FBS, RMBIO) in a humidified atmosphere with 5% CO_2_ at 37°C. THP-1 cells were differentiated to macrophages by supplementing the growth medium with 200 nM phorbol myristate acetate (PMA, Sigma) overnight.

### Plasmid construction

The pBESTluc plasmid was used as a backbone for all reporter assay constructs and was included in the Luciferase Assay System kit (Promega). pBEST + *metK* was created by inserting nucleotides corresponding to the first 10 codons of the *C. burnetii metK* gene immediately downstream of the *luc* start codon using a Q5 Site-Directed Mutagenesis Kit, as instructed (New England Biolabs). *cbsR12* was cloned into pBESTluc using specific primers containing XhoI and AfeI restriction sites on the forward and reverse primers, respectively. The forward primer also encoded a P*tac* promoter and *lac* operator. The resulting PCR product was cloned into pBEST + *metK* using unique XhoI and AfeI restriction sites in an irrelevant intergenic region. A frameshifted *luc* construct was created as a byproduct of the *metK* Q5 mutagenesis of the pBESTluc plasmid. This construct contained a 1-bp frameshift deletion in the 5’ end of *luc*. PBEST + *carA* 5’ UTR was created using primers specific to the 5’ UTR of *carA* with HindIII and BamHI restriction sites on the forward and reverse primers, respectively. The forward primer also encoded the P*tac* promoter. Nucleotides corresponding to the *lac* operator were inserted using a Q5 Site-Directed Mutagenesis Kit to create the final pBEST + *carA* 5’ UTR construct. The *cbsR12* gene was inserted into this construct in the same fashion as for pBEST + *metK* + *cbsR12*.

Recombinant CarA and MetK were generated by PCR amplification of *carA* and *metK* using forward and reverse primers containing BamHI and HindIII restriction sites, respectively. The resulting amplicons were cloned into compatible restriction sites of pQE30 (Qiagen) by standard protocol.

### Axenic growth of *C. burnetii*

For growth curves and Crosslink-Seq experiments, 10^6^ GE/mL of either *Tn1832*, *Tn327*, or *Tn327-Com*p were inoculated into 300 mL ACCM-2 in a 1-liter flask with either chloramphenicol (5 μg/mL), kanamycin (350 μg/mL) or both. GE/mL was initially determined from frozen cell stocks by quantitative PCR analysis as previously described [76], although different *dotA* primers were used (**S9 Fig**). Cell viability was then determined using a BacLight Bacterial Viability kit as instructed (Thermo Scientific).

### *C. burnetii* infection of differentiated THP-1 cells

THP-1 cells were seeded onto 4-well chambered glass slides (Labtek) or T-75 flasks. After two days growth, 200nM PMA was added along with fresh medium and the cells were allowed to differentiate overnight. The PMA-supplemented medium was removed and fresh medium was restored, after which differentiated THP-1s were allowed to recover for 4 hours prior to *C. burnetii* infection at an MOI of 10. Initial infections were rocked for 2 hours at room temperature before returning cells to 5% CO_2_ and 37°C. At 1dpi the supernatant was removed, extracellular *C. burnetii* was washed away with warmed 1X PBS, and fresh medium was added.

### Total RNA and genomic DNA extraction and purification

*C. burnetii* grown in ACCM-2 was centrifuged at 15,000 x g at 4°C for 15 minutes, after which pellets were resuspended in 1 mL TRI Reagent (Ambion). The suspension was incubated for 1 hour at room temperature, frozen for 2 hours at −80°C, thawed for 30 minutes at room temperature, then pipetted vigorously until homogenized. 100 µL BCP (Acros Organics) was added, the solution was vortexed for 30 seconds, incubated for 5 minutes at room temperature, and centrifuged at 12,000 x g at 4°C for 10 minutes. The aqueous phase was then collected and 300 µL 100% ethanol added. The mixture was immediately vortexed for 10 seconds, and a RiboPure RNA Purification kit (Ambion) was used to collect, concentrate, and wash the resulting total RNA. The RNA was collected in nuclease-free H_2_O and treated with DNase I for 1 hour at 37°C. After DNase inactivation, the total RNA was precipitated for 3 hours at −80°C in a mixture of glycogen, ammonium acetate and 100% ethanol, and the precipitate collected by centrifugation for 30 minutes at 16,000 x g. The resulting purified RNA was run on a NanoDrop spectrophotometer (Thermo Scientific) to determine concentration and purity.

In order to purify total RNA from *C. burnetii* grown in THP-1 cell lines, growth medium was first removed and replaced with 1 mL TRI Reagent. Flasks containing TRI Reagent were then rocked for 1 hour at room temperature, after which cells were mechanically scraped and collected into a 15-mL conical tube. The mixture was frozen overnight at −80°C, thawed to room temperature for 30 minutes, and the RNA purification procedure continued as described above.

Genomic DNA was purified from TRI Reagent mixtures according to manufacturer protocols (Ambion). Briefly, after the aqueous phase was removed in the total RNA extraction, the resulting mixture was centrifuged at 12 x g for 10 minutes, the supernatant (protein) removed, the pellet washed twice with a DNA wash solution (0.1M trisodium citrate, 10% ethanol) followed by 75% ethanol. The resulting pellet was air-dried and then solubilized overnight into 8mM NaOH at 4°C. The mixture was centrifuged and the supernatant, containing genomic DNA, was collected. 40 µL 0.1M HEPES, free acid (pH ∼5.5) was then added and the resulting DNA was purified using a Nucleotide Removal kit as instructed (Qiagen).

### Quantitative PCR (qPCR) and quantitative real-time PCR (qRT-PCR)

qPCR and qRT-PCR experiments were performed as previously described [77]. Briefly, 1 µg total RNA was subjected to random-primed cDNA synthesis using an iScript cDNA Synthesis kit following the manufacturer’s protocol (Bio-Rad). 1 µL of the resulting cDNA was used in triplicate 25-µL qRT-PCR reactions, including 300 nM each of primers specific to *cbsR12* and a volume of iQ SYBR Green Supermix (Bio-Rad). The resulting reactions were cycled on a MyiQ Single-Color Real Time PCR Detection System (version 1.0 software) (Bio-Rad). CbsR12 cDNA copy number was normalized against *dotA* copy number derived from *C. burnetii* genomic DNA to obtain copy numbers / GE values. Growth curve copy numbers were obtained from genomic DNA purified from the same cells from which total RNA was purified.

### RNase III assay

RNase III Assay was performed as previously described [46]. Briefly, 200 nM substrate RNA was incubated for 30 minutes at 37°C with or without recombinant *C. burnetii* RNase III or *E. coli* RNase III. *C. burnetii* intervening sequence (IVS) RNA was used as a positive control substrate [46]. The resulting reactions were electrophoresed on a 7% denaturing polyacrylamide gels stained with 2 μg/mL acridine orange to visualize.

### Identification of transcription start sites

5’ RACE analysis of *carA*, *metK*, *cvpD*, and *cbsR12* transcripts was performed on *C. burnetii* total RNA extracted from LCVs (3dpi) infecting Vero (*carA* and *metK*) or THP-1 (*cvpD*) cells using a 5’ RACE System kit (Invitrogen) according to manufacturer protocols and with gene-specific primers (see **S9 Fig**). Resulting PCR products were cloned into pCR2.1-TOPO as instructed (Invitrogen) and then sequenced with M13 universal primers by Sanger automated sequencing.

### *In silico* and bioinformatics analyses

RNA target predictions were carried out using TargetRNA2 [78], IntaRNA [79], and CopraRNA [80] algorithms and default settings. RNA was folded using mFold [81] and visualized with the Visualization Applet for RNA [82]. Analyses of RNA-Seq data were carried out as previously described [31]. Briefly, raw fastq files were concatenated, quality-filtered with the FASTX toolkit (http://hannonlab.cshl.edu/fastx_toolkit/) and then clipped, aligned, and filtered with Nesoni version 0.128 tools (http://www.vicbioinformatics.com/software.nesoni.shtml). Transcripts per million (TPM) were calculated using custom perl and python scripts that can be accessed through GitHub (https://github.com/shawachter/TPM_Scripts). Crosslink-Seq enrichment was accomplished by processing .bam files using featureCounts [83], followed by use of the DESeq2 package in R version 3.4.4 to obtain differentially expressed genes [84]. The Artemis genome browser was used to visualize generated alignment files (http://www.sanger.ac.uk/science/tools/artemis) [85].

All IFA images were processed and analyzed using Fiji [86] and Cell Profiler [87], respectively. Figures were created using R version 3.4.4, Inkscape (https://inkscape.org/release/inkscape-0.92.4/) and GIMP (https://www.gimp.org/downloads/).

### RNA-RNA hybridization and electrophoretic mobility shift assay

PCR products (1μg) of desired templates were *in vitro-*transcribed overnight at 37°C with 2.5 mM Ribonucleotide Solution Mix (New England Biolabs) and when needed, 0.5 mM Bio-16-UTP (Invitrogen) using a MAXIscript T7 Transcription kit (Invitrogen). Resulting reactions were incubated for 1 hour at 37°C with 1 µL TURBO DNase (Invitrogen) after which 21 µL 2X formaldehyde loading dye (Thermo Scientific) was added. Reactions were heated for 4 minutes at 85°C then immediately plunged on ice. Samples were electrophoresed on a 7% polyacrylamide gel for 75 minutes at 100V, and the gel was stained with a 2 μg/mL acridine orange solution for 10 minutes and de-stained with milli-Q water for 30 minutes. Visualized bands were excised and eluted overnight into probe elution buffer (0.5M AmAc, 1mM EDTA, 0.2% SDS) at 37°C. The resulting solution was precipitated with ethanol overnight at −20 °C, washed with 70% ethanol, and resuspended in nuclease-free H_2_O. RNA concentrations were determined using a NanoDrop spectrophotometer. Following this, 10 nM biotin-labeled CbsR12 and an equal concentration of target RNA were combined and heated for 5 minutes at 85°C. A high-salt TMN buffer (100 mM NaCl, 50 mM MgCl_2_, 100 mM Tris-Cl, 0.05% Tween-20) was then added and the reactions were immediately plunged on ice for 30 seconds and then incubated for 30 minutes at 37°C. A non-denaturing loading dye (0.25% Bromophenol blue) was added and the resulting RNA mixtures were resolved on a 7% polyacrylamide gel for 1 hour 20 minutes at 100V. RNA was transferred to a BrightStar-Plus Positively Charged Nylon Membrane (Ambion) using an electro-blot transfer system (Bio-Rad) and cross-linked with short-wave UV light in a GS Gene Linker UV Chamber (Bio-Rad). A North2South Chemiluminescent Hybridization and Detection Kit (Thermo Scientific) was used to detect the resulting bands. The blot was imaged on a LAS-3000 imaging system (Fujifilm).

### RNA-protein electrophoretic mobility shift assay

The CbsR12-CsrA1/2 EMSA was performed as previously described for CsrA-binding RNAs [47]. Briefly, biotin-labeled CbsR12 was synthesized *in vitro* as described above for RNA-RNA EMSAs. *C. burnetii csrA-1* and *csrA-2* genes were cloned into the expression vector pQE30, expressed, and natively purified as previously described [46]. 1 nM biotin-labeled CbsR12 diluted in TE buffer (10mM Tris-HCL, 1mM EDTA) was heated at 75°C for 3 minutes and equilibrated to room temperature for 10 minutes. Purified CsrA1/2 diluted in CsrA-binding buffer (1 µl in 10mM Tris-HCl, 10mM MgCl_2_, 100mM KCl, 10mM DTT, 10% glycerol, and 10 U RNasin [Promega]) was added, and reactions were incubated for 30 minutes at 37°C. The samples were immediately resolved on a 10% non-denaturing polyacrylamide gel for 3h at 125V, transferred to a BrightStar-Plus Positively Charged Nylon Membrane (Ambion) using an electro-blot transfer system (Bio-Rad), and cross-linked with short-wave UV light in a GS Gene Linker UV Chamber (Bio-Rad). A North2South Chemiluminescent Hybridization and Detection Kit (Thermo Scientific) was used to detect the resulting bands. The blot was imaged on a LAS-3000 imaging system (Fujifilm). The K_D_ for CsrA-2 was determined as previously described [47].

### Reporter assay

A Luciferase Assay System kit (Promega) was used. All pBESTluc reporter assay constructs were transformed into *E. coli* Top 10 F’. Resulting *E. coli* strains were grown overnight at 30°C in 10 mL LB containing ampicillin (100 µg/mL) and 1% glucose. An aliquot (4.5 mL) of the overnight culture was inoculated into 40.5 mL LB with 100 µg/mL ampicillin and grown for 1.5 hours at 30°C. IPTG was added (to 1 mM) and culture aliquots (100 μl) were removed at 0, 1, and 2 hour time points. 80 µL LB and 20 µL CCLR lysis solution (1X CCLR, 25 mg BSA, 12.5 mg lysozyme, 7.5 mL water) were added to the aliquots and gently inverted until the solution clarified. 50 µL of the resulting lysate was aliquoted to a 96-well plate, 100 µL of luciferase assay substrate was added, and luminescence was immediately read with a SpectraMax M5 Plate Reader (Molecular Devices).

### Crosslink-Seq analysis

*In vivo* RNA-RNA crosslinking was performed as previously described with minor modifications [50, 88]. *C. burnetii Tn1832* or *Tn327* LCVs from axenic culture were collected by centrifugation at 15,000 x g at 4 °C for 15 minutes, washed twice in 1X PBS (pH 7.4), and resuspended in 5 mL fresh 1X PBS. The cells were incubated for 10 minutes on ice and 0.2 mg/mL 4’-aminomethyltrioxsalen (AMT, Cayman Chemicals) was added and mixed. This was followed by exposure to long-wave (365 nm) UV light for 1 hour on ice. The cells were centrifuged as above, briefly washed with 1X PBS and re-centrifuged. The cellular pellet was resuspended in 1 mL TRI Reagent and total RNA was extracted. The RNA was DNase I treated and then mixed with an equal volume of hybridization buffer (20 nM HEPES pH8, 5 mM MgCl2, 300 mM KCl, 0.01% NP-40, 1 mM DTT). This mixture was heated for 5 minutes at 85°C then cooled on ice. 10 nmol of two distinct biotinylated anti-CbsR12 probes were added to the mixture and incubated overnight at room temperature. 150 µL of NeutrAvidin resin (Thermo Scientific) was washed twice in WB100 buffer (20 mM HEPES pH 8, 10 mM MgCl2, 100 mM KCl, 0.01% NP-40, 1 mM DTT) then blocked for 2 hours in blocking buffer (WB100, 50 µL BSA (10 mg/mL), 40 µL tRNA (10 mg/ mL), 10 µL glycogen (20 mg/mL)). The total RNA mixture hybridized to the biotinylated anti-CbsR12 RNAs were mixed with the resin and incubated for 4 hours at 4°C. The resin was then washed 5 times in WB400 buffer (20 mM HEPES pH 8, 10 mM MgCl2, 400 mM KCl, 0.01% NP40, 1 mM DTT) and the hybridized RNAs were re-isolated with TRI Reagent. The remaining RNAs were un-crosslinked by exposure to short-wave UV light (254 nm) for 15 minutes on ice. The resulting RNA was sent to the Yale Center for Genomic Analysis for RNA-Seq. The sequencing reads are available at the NCBI sequencing read archive (accession number: SUB5191080).

### Immunofluorescence assay

IFAs on infected THP-1 cells were performed as previously described with modifications [14]. Briefly, 4-well chambered glass slides were coated for 30 minutes with a 0.2% solution of Sigmacote (Sigma). THP-1 cells were inoculated into the chambered slides and incubated overnight or until 60% confluence was reached. Confluent cells were then infected with *Tn1832*, *Tn327*, or *Tn327-Comp C. burnetii* strains at a MOI of 10. At 1dpi, the infection was stopped by washing cells three times for 5 minutes in pre-warmed 1X PBS, after which fresh medium was re-added. At 3 or 7 dpi, the growth medium was removed and cells were fixed with ice-cold 100% methanol for 5 minutes at room temperature. Cells were washed three times for 5 minutes each with ice-cold 1X PBS, blocked for 1 hour at room temperature with a 2% BSA solution in 1X PBS, and then incubated with anti-Com1 (1:1000) and anti-LAMP1 (1:50, H4A3 was deposited to the Developmental Studies Hybridoma Bank by August, J.T / Hildreth, J.E.K) antibodies for 2 hours. Cells were washed and incubated with Alexa Fluor 488 (1:200, Thermo Scientific) and goat anti-mouse rhodamine antibodies (1:200, Thermo Scientific) along with DAPI (300 nM, Thermo Scientific) for 1 hour. Cells were then washed three times for 5 minutes each in ice-cold 1X PBS and immediately imaged. Images were processed with Fiji [86]. Cell Profiler was used to count colonies per cell and measure CCV areas, as previously described [89].

### Protein expression, purification and antibody production

Recombinant *Coxiella* RNase III was expressed from a previously generated pQE30 construct and purified as before [46]. *C. burnetii carA* and *metK* genes were cloned in frame into the pQE30 expression vector (Qiagen) and the resulting N-terminal His_6_-tagged proteins were expressed and purified as previously described for the *C. burnetii* RNA helicase [90]. Purified recombinant CarA and MetK proteins were submitted to General Bioscience, Inc., for rabbit polyclonal antibody production.

### Western blot

Western blots were performed as previously described [29]. Briefly, *Tn1832*, and *Tn327-Comp* were grown for 4 days (6 days for *Tn327*) in ACCM-2. Bacterial proteins were prepared by lysing and solubilizing pellets into Laemmli sample buffer (50 mM Tris-HCl, 4% (w/v) SDS, 10% (v/v) glycerol, 0.1% (w/v) bromophenol blue, 5% (v/v) β-mercaptoethanol) followed by boiling for 10 min. Proteins at 30 µg (CarA blot) or 60 µg (MetK blot) were resolved on 10% - 20% acrylamide gradient Tris-Glycine SDS-PAGE gels. Gels were immediately electro-transferred to a nitrocellulose membrane for 1 hour at 100V in transfer buffer (25 mM Tris, 192 mM glycine, 20% v/v methanol, pH 8.3). The resulting blot was blocked (1X PBS pH 7.4, 0.3% (v/v) Tween-20, 2% (w/v) non-fat milk) for 1 hour, then incubated with primary antibody solution (1X PBS pH7.4, 0.3% (v/v) Tween-20, 1:500 CarA / 1:5000 MetK primary antibody) for 2 hours with rocking at room temperature. The blot was then washed five times for 5 min in 1X PBS, then incubated for 1 hour in secondary antibody solution (1X PBS + 1:2000 goat anti-rabbit::HRP). The blot was again washed five times for 5 minutes in 1X PBS and immediately developed using SuperSignal West Pico Chemiluminescent Substrate (Thermo Scientific) according to the manufacturer’s protocol. Imaging was performed on a ChemiDoc XRS+ System (Bio-Rad).

### Generation of a CbsR12 complemented strain

The *cbsR12* mutant (strain *Tn327*) was complemented as previously described, with modifications [91]. Briefly, the *cbsR12* gene along with 100 bp of flanking sequences were PCR-amplified using primers containing EcoRI and BamHI restriction sites. The amplicon was cloned into compatible restriction sites of the pMini-Tn7-KAN plasmid by standard protocol [92]. The resulting plasmid was transformed into electrocompetent *E. coli* PIR1 cells for propagation. The pMini-Tn7-CbsR12-KAN plasmid (20 μg), along with 10 μg of a second plasmid containing the transposase, pMini-TnS2-ABCD, were transformed into *C. burnetii Tn327* in a single electroporation reaction (25 kV, 500 ohms, 25 μF). Electroporated cells were allowed to recover for 5 days in ACCM-2 supplemented with 1% FBS, and dilutions were plated onto ACCM-2 agar containing kanamycin (375 μg/mL). Isolated colonies were picked and re-cultured on ACCM-2 agar plates for several rounds. Colony-PCR was used to screen for the *Tn327-Comp* strain, and the location of the *cbsR12* cassette was determined by PCR and Sanger automated sequencing. qPCR of *Tn327-Comp* genomic DNA utilizing primers specific to *cbsR12* was used to ensure that a single transposon insertion event occurred.

## Acknowledgments

We would like to thank Paul Beare for his generous donation of *E. coli* PIR1 cells, pMini-TnS2-ABCD plasmid, and the pMini-Tn7-KAN plasmid. We would also like to thank Jenny Wachter for her contribution of the TPM calculator and Linda D. Hicks for her excellent technical assistance.

## Supporting information

**Fig S1. CbsR12 is processed by RNase III.** (**A**). CbsR12 secondary structure as predicted by mFold. Nucleotide 1 was determined to be the TSS for the full-size transcript by 5’ RACE. Red asterisks indicate apparent alternative TSSs from 5’ RACE analysis. The dotted line indicates the putative RNase III processing area. The blue solid lines indicate consensus CsrA-binding sites. (**B**). RNase III assay of *in vitro-*transcribed CbsR12 with the *C. burnetii* IVS RNA as a positive control substrate. Results from treatment with *E. coli* (*Ec*) RNase III (New England BioLabs), recombinant *C. burnetii* (*Cb*) RNase III or no-enzyme controls are shown and were done as previously described [46]. Arrows indicate RNase III-processed (blue) and un-processed (red) CbsR12 RNA.

**Fig S2. Location of the *Tn327* transposon insertion to inactivate *cbsr12*.** (**A**). The *cbsR12* gene and promoter elements are highlighted in various colors, while the location of the *Himar* transposon insertion producing the *Tn327* strain of *C. burnetii* is indicated by a black arrow. Red arrows indicate primer-binding sites for PCR confirmation of the lesion (forward and reverse primers above and below their annealing sequences, respectively). (**B**). PCR products confirming transposon insertion in the *cbsR12* gene of *Tn327* (by loss of the ∼250 bp amplicon) and reintroduction of *cbsR12* in *Tn327-Comp*.

**Fig S3. CbsR12 competitively binds *carA* transcripts in a dose-dependent manner.** RNA-RNA EMSA showing hybridization reactions between biotin-labeled CbsR12 (Bio-CbsR12) and an *in vitro-*transcribed segment of *carA*. Anti-CbsR12 represents a positive control consisting of a transcript equal in size but antisense to the CbsR12 transcript. A cold-chase sample containing Bio-CbsR12 + un-labeled CbsR12 + CarA shows competitive (specific) binding relative to Bio-CbsR12 + CarA, while increasing the dose of *carA* transcript (from 2 nM to 10 nM) increases the amount of retarded sample signal on the blot. Arrows indicate un-bound Bio-CbsR12 (blue) and Bio-CbsR12 bound to its RNA targets (red).

**Fig S4. CbsR12 competitively binds *metK* transcripts in a dose-dependent manner.** RNA-RNA EMSA showing hybridization reactions between biotin-labeled CbsR12 (Bio-CbsR12) and an *in vitro*-transcribed segment of *metK*. Anti-CbsR12 represents a positive control consisting of a transcript equal in size but antisense to the CbsR12 transcript. A cold-chase sample containing Bio-CbsR12 + un-labeled CbsR12 + MetK shows competitive (specific) binding relative to the signal produced in Bio-CbsR12 + MetK, while increasing the dose of *metK* transcript (from 2 nM to 10 nM) increases the amount of retarded sample signal seen on the blot. Arrows indicate un-bound bio-CbsR12 (blue) and bio-CbsR12 bound to its RNA targets (red).

**Fig S5. CbsR12 competitively binds *cvpD* transcripts in a dose-dependent manner.** RNA-RNA EMSA showing hybridization reactions between biotin-labeled CbsR12 (Bio-CbsR12) and an *in vitro-*transcribed segment of *cvpD*. Anti-CbsR12 represents a positive control consisting of a transcript equal in size but antisense to the CbsR12 transcript. A cold-chase sample containing Bio-CbsR12 + un-labeled CbsR12 + CvpD shows competitive (specific) binding relative to the signal produced in Bio-CbsR12 + CvpD, while increasing the dose of *cvpD* transcript (from 2 nM to 10 nM) increases the amount of retarded sample signal seen on the blot. Arrows indicate un-bound bio-CbsR12 (blue) and bio-CbsR12 bound to its RNA targets (red).

**Fig S6. CbsR12 binding to *metK* and *ahcY* transcripts are separate events.** Artemis representation of Crosslink-Seq results for *Tn1832*. Red and blue lines represent the two biological replicates. The blue and red arrows indicate *metK* and *ahcY* reads, respectively, that crosslinked with CbsR12.

**Fig S7. CbsR12 downregulates the quantity of transcripts arising from the 5’ end of *cvpD* in LCVs from infected THP-1 cells.** (**A**). *cvpD* gene sequence from the 5’ TSS to the predicted downstream alternative start codon. Various colors highlight the TSSs, the CbsR12-binding site, the putative downstream promoter, putative RBSs, and start codons. Red arrows indicate primer annealing regions for qRT-PCR (forward and reverse primers above and below their respective annealing sequences). (**B**). qRT-PCR of the 5’ end of *cvpD* from *Tn1832*, *Tn327*, and *Tn327-Comp* LCVs (3dpi) and SCVs (7dpi) infecting THP-1 cells. Values represent the means ± standard error of means (SEM) of three independent determinations (** = P < 0.01, one-way ANOVA, *** = P < 0.001, one-way ANOVA).

**Fig S8. CbsR12 induction in *E. coli* leads to an autoaggregative phenotype.** Overnight cultures of *E. coli* Top10 F’ harboring pBEST + *carA*5’UTR or pBEST + *carA*5’UTR + *cbsR12* were passaged into 3 mL fresh LB supplemented with ampicillin (100 µg/mL) for 2 hours, then induced with 1 mM IPTG for 3 hours before photography. The red arrow indicates autoaggregation of *E. coli* upon CbsR12 induction.

**Fig S9. Strains, plasmids, and primers used in the study.**

